# The Chromatin Protein CFDP1 Activates TPX2 and Promotes Chromosomal Microtubule Nucleation and Spindle Assembly

**DOI:** 10.1101/2025.09.22.677689

**Authors:** Gokul Gopinathan, Xianghong Luan, Thomas G.H. Diekwisch

**Affiliations:** School of Medicine and Dentistry, University of Rochester, Rochester, NY

**Keywords:** CFDP1, TPX2, Microtubule nucleation, K-fibers, spindle

## Abstract

Microtubule associated proteins (MAPs) are multifunctional tubulin-binding proteins that contribute to essential aspects of mitotic spindle formation. In the present study, loss of the MAP CFDP1 in mice resulted in gastrulation defects and embryonic lethality at e8.5 due to chromosome segregation spindle defects and loss of K-fiber stability. CFDP1 decreased the association of the nuclear transport protein importin α with the essential spindle assembly factor TPX2, thereby promoting Aurora A kinase activation, microtubule nucleation and spindle assembly. Further defining CFDP1 mode of action we identified CFDP1 as a bipartite molecule with an acidic N-terminus that harbors a nuclear localization signal essential for importin α dissociation from TPX2 and a basic C-terminus that interacts with tubulin, co-localizes with the mitotic spindle, and promotes microtubule bundling and polymerization. Together, our studies have established CFDP1 as an essential bipartite MAP involved in chromosomal microtubule nucleation in conjunction with TPX2 and Aurora A while also facilitating nuclear TPX2 activation through importin α dissociation.

## INTRODUCTION

Assembly of a microtubule (MT)-based spindle apparatus that faithfully partitions genetic information during cell division is a prerequisite for the viability of eukaryotic organisms. In animal cells, the centrosomes are major nucleators of microtubules which are subsequently organized into a bipolar spindle.^[1]^ However, several non-centrosomal pathways of MT nucleation and spindle assembly have been documented in cells and developing embryos which are normally masked by the presence of the centrosome.^[2,3]^ For example, during mitosis MTs are nucleated near chromosomes in a process that depends on the activity of the small GTPase, Ran.^[4,5]^ Chromatin-induced microtubule assembly functions through the Ran-GTP pathway in which RCC1 mediated generation of a Ran-GTP gradient in the immediate vicinity of chromosomes facilitates MT nucleation through local dissociation of several spindle assembly factors (SAFs) including TPX2.^[6–8]^ These non-centrosomal MTs are captured, stabilized and assembled into ordered arrays by several proteins including microtubule associated proteins (MAPs) by virtue of their ability to modulate MT physical properties. Ultimately, non-centrosomal MTs are combined with the MTs originating from the centrosomes, seamlessly integrating them into a common spindle structure.^[9]^ Underscoring their importance during mitosis, chromosome-mediated MT assembly is demonstrated to be a significant contributor to mitotic spindle assembly even in cells containing centrosomes.^[10]^ Mitotic spindle assembly and its link to sister chromatids during mitosis occurs at a disc-shaped protein structure called kinetochore.

Kinetochores are macromolecular protein machines assembled at the centromere that mediate chromosome segregation by linking chromosomes to spindle microtubules.^[11]^ The mitotic spindle fibers that join kinetochores to the spindle poles are called K-fibers.^[12]^ The K-fiber spindle microtubules (MTs) are attached at the kinetochores of corresponding chromosomal centromeres, where motor proteins generate forces to power chromosome movement toward daughter nuclei.^[11,12]^ K-fiber formation occurs through two distinct mechanisms.^[13]^ The first mechanism involves the capture of an astral MT produced at the centrosome by a kinetochore and was directly visualized in vertebrate cells.^[14,15]^ In the second mechanism, short microtubules nucleated at the kinetochores are captured and oriented by CENP-E/kinesin-7 motors, ^[16]^ which are then incorporated into the spindle by cytoplasmic dynein motors which transport the growing fiber pole-ward along non-kinetochore microtubules.^[17]^ More recently, branching microtubule nucleation has been discovered as an additional mechanism in which MT in K-fibers are nucleated from pre-existing spindle MTs in an augmin-dependent manner. ^[18,19]^ Although the relative contribution of these mechanisms toward normal spindle assembly is not clear, all three mechanisms are known to coexist and aid in mitotic progression. Non-centrosomal MTs nucleated at the kinetochores present a significant advantage as these can be easily captured and stabilized by the kinetochore complex favoring the “search-and-capture” model of kinetochore-MT attachment and chromosome segregation.^[20]^ In support, several studies in somatic cells have demonstrated that non-centrosomal MTs form primarily in the vicinity of the centromere and not around chromosome arms.^[17,21,22]^

Microtubule assembly and dynamics are precisely regulated in time and space by microtubule-associated proteins (MAPs), a group of proteins which are involved in microtubule nucleation, stabilization, and transport of cargo along microtubules. Modulation of MT mechanical properties by MAPs is a key process that controls MT-MT encounters resulting in formation of complex MT bundle arrays in eukaryotic cells.^[23]^ MT deformations induced by MAPs primarily alter MT flexibility and have been suggested to be essential for the generation of MT bundles.^[24,25]^ The neuronal MAPs tau and MAP2 are known to stiffen MTs by decreasing MT flexular rigidity *in vitro* ^[26]^ whereas MAP65-1/Ase1, from the conserved MAP65 family drastically increases MT flexibility resulting in a softening effect on MTs.^[23]^ MT softening by MAPs was suggested as a general mechanism regulating MT network plasticity and MT bundling.^[23]^ Several MAPs including kinesin-12 Kif15, TPX2, clathrin/chTOG/TACC3 complex, HURP, and kinesin Kif18A preferentially localize to K-fibers and stabilize them enabling a reliable attachment between kinetochores and the spindle poles.^[27–31]^ In addition, a large number of chromatin proteins have been shown to regulate MT dynamics during mitosis.^[32]^ Chromatin associated MAPs such as Dppa2, kinesin-4/KIF4 and NuSAP bind chromatin and MTs simultaneously *via* distinct chromatin and MT binding domains, whereas chromatin-dissociated MAPs including, CHD4, INO80, ISWI and KANSL dissociate from chromatin upon mitotic entry to perform MT related functions *via* their chromatin-binding nuclear localization signal (NLS) region.^[32,33]^

In earlier studies, CFDP1 (Craniofacial development protein 1) was identified as an essential protein involved in cell proliferation, survival of mouse fibroblasts, and craniofacial development.^[34–36]^ The Drosophila homolog, *Yeti* was essential for larval development as a result of its role in higher-order chromatin organization,^[37]^ whereas the yeast homolog, *Swc5* functions within the SWR1 chromatin remodeling complex to facilitate H2A.Z histone variant exchange.^[38]^ Interestingly, Yeti was originally defined as a kinesin binding protein able to bind both subunits of the microtubule-based motor kinesin-I.^[39]^ Furthermore, a proteomics study of mitotic chromosome composition in chicken cells localized CFDP1 (Centromere Protein 29, CENP-29) to the outer kinetochore region.^[40]^ In the present study we have hypothesized that CFDP1 functions as a microtubule associated protein (MAP) based on its role in microtubule nucleation, stabilization and elongation. Based on this hypothesis we sought to localize CFDP1 in relationship to the mitotic spindle, determine its effect on spindle integrity, cell proliferation and embryonic development, and decipher the contributions of its functional domains toward TPX2 mediated tubulin nucleation and Aurora A kinase activation.

## RESULTS

### Gastrulation defects and early embryonic lethality in *Cfdp1* null mice

To determine CFDP1 function during mammalian development, a knockout (KO) mouse model was generated replacing the first exon with a LacZ-Neomycin cassette (Figure S1A and S1B, Supporting Information). Mating between *Cfdp1* knockout heterozygotes (*Cfdp1*^+/-^ X *Cfdp1*^+/-^) yielded wild-type (WT) and heterozygous offspring at Mendelian frequencies while homozygous null mice (*Cfdp1*^-/-^) were lost (63-live births, 21-WT, 42-heterozygotes, 0-KO) (Figure S1C, Supporting Information). Detailed analysis of embryos derived from *Cfdp1*^+/-^ intercrosses indicated that KO embryos (*Cfdp1*^-/-^) were not viable at embryonic day 8.5 (e8.5) and were resorbed by e12.0, suggesting lethality during gastrulation and early organogenesis (Figure S1C, Supporting Information).

Immunohistochemical analysis using anti-CFDP1 antibody revealed a complete absence of CFDP1 protein in e3.5 KO blastocysts (Figure 1A and 1B). We then performed blastocyst outgrowth experiments in ES cell media devoid of LIF (Leukemia inhibitory factor) resulting in trophoblast ectodermal cellular outgrowth. While the wild-type group featured distinct inner cell mass (ICM) condensations, the ICM was lost in *Cfdp1* KO blastocysts and only a few single cells remained (Figure 1C). At e7.5, *Cfdp1* KO embryos were significantly smaller and displayed rudimentary embryonic tissues lacking germ layer organization when compared to same stage mid-gastrulation wild-type embryos featuring well differentiated germ layers (Figure 1D). Analysis of cell proliferation and tissue organization in *Cfdp1* KO embryos revealed significantly reduced Bromodeoxyuridine (BrdU) incorporation (Figure 1E) and decreased laminin expression (Figure 1F). Furthermore, intraembryonic tissues in e6.5 *Cfdp1*^-/-^ embryos were compressed, and the conceptus was predominantly populated by a condensation of GATA4 positive extraembryonic endoderm (Figure 1G). There was no distinct epiblast in *Cfdp1*^-/-^ embryos, only a rudimentary extraembryonic ectoderm layer labeled by the ectoderm marker EED remained (Figure 1H). Remarkably, absence of SNAIL immunoreactivity indicated a lack of mesoderm in *Cfdp1* KO embryos at e6.5 when compared to WT controls (Figure 1I). These data demonstrated that *Cfdp1* is essential for gastrulation and early organogenesis in mice, severely impacting development.

**Figure 1.**
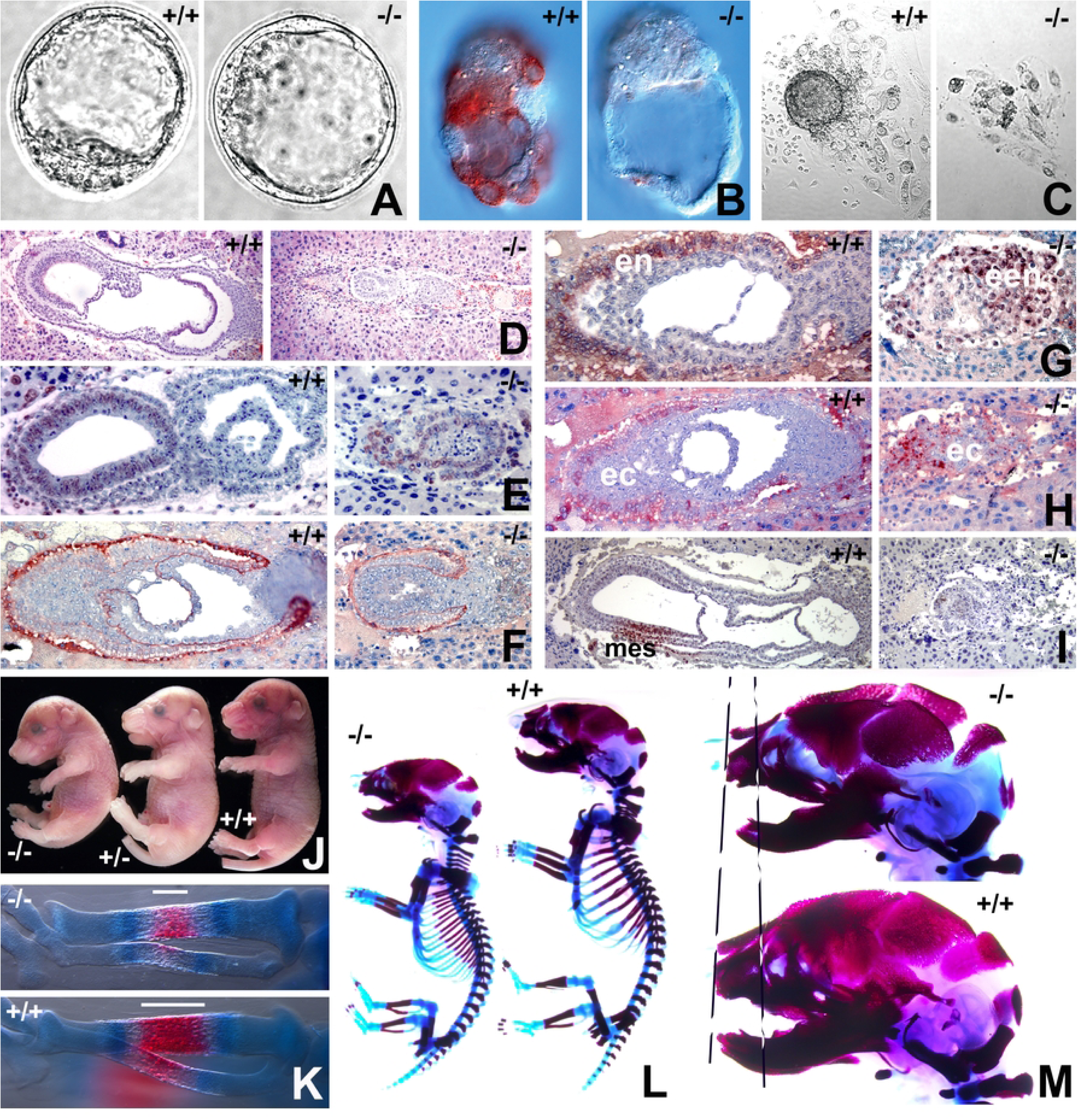
Mid gastrulation defects and developmental delays in mice lacking *Cfdp1*. A-I) *Cfdp1* null phenotype in mouse blastocysts and embryos. A) *Cfdp1^-/-^* and wild-type blastocysts at e3.5. B) CFDP1 localization in *Cfdp1^-/-^* and wild-type blastocysts. C) Blastocyst outgrowth and inner cell mass (ICM) condensations in e3.5 wild-type and *Cfdp1^-/-^* blastocysts. D) Rudimentary *Cfdp1^-/-^* embryonic tissues (e7.5) compared to same-stage wild-type embryos featuring ectoderm (ect), mesoderm (mes), and endoderm (end) germ layers. E) Reduced BrdU incorporation in *Cfdp1^-/-^* embryos compared to controls. F) Laminin expression in wild-type and *Cfdp1^-/-^* embryos. G) A major portion of *Cfdp1^-/-^* embryonic tissue was stained with the extraembryonic endoderm tissue marker GATA4. H) Diminished and disorganized extraembryonic ectoderm (EED) in *Cfdp1^-/-^* embryos. I) Absence of the mesoderm marker SNAIL in *Cfdp1^-/-^* embryos. J) Developmental delays in *Cre* mediated conditional *Cfdp1* deletion mice (-/- homozygous and +/- heterozygous). Decreased mineralization in (K) tibia and L) skeletal tissues of *Cfdp1* conditional knock-out embryos (e16.5). (M) Craniofacial developmental defects in *Cfdp1* conditional knock-out embryos (-/-) compared to wild-type littermates (+/+). Representative skeletal preps were imaged in K, L and M.

To circumvent the early embryonic - lethal phenotype, we next generated mice harboring conditional knock out alleles for *Cfdp1*. These floxed mice were generated by inserting a LoxP/FRT flanked Neomycin cassette spanning exon1 and the flanking 3’ region of *Cfdp1* (*Cfdp1*^flox/flox^) (Figure S1D and S1E, Supporting Information). For conditional knock out of *Cfdp1*, mice harboring a floxed allele and a KO allele for *Cfdp1* were mated to *Rosa26 Cre* mice and recombination was initiated in resulting test (Rosa26Cre; Cfdp1^-/flox^) and control mice (Rosa26Cre; Cfdp1^+/flox^) by Tamoxifen administration. Since conditional deletion of *Cfdp1* in adult test mice did not result in an apparent phenotype (data not shown), we initiated *Cfdp1* deletion *in utero* in embryos by Tamoxifen (2 mg i.p) administration to pregnant mice (at e11.5 gestation). Confirming the essential role of CFDP1 during early mouse development, we did not obtain any live offspring post tamoxifen administration in the inducible knockout model and on several occasions *Cfdp1* conditional KO embryos were born dead (data not shown).

*Cfdp1* conditional deletion in embryos at a later developmental stage (tamoxifen administration at e13.5 and harvested at e18.5) resulted in embryos which were significantly smaller in size and exhibited a distinct curvature of the spine compared to heterozygous and wild type littermates (Figure 1J). Whole mount staining of e18.5 embryos with Alizarin Red and Alcian Blue revealed a shorter embryonic skeleton (∼14.9% shorter) and an overall decrease in bone mineralization among the *Cfdp1* conditional KO embryos (Figure 1, K-M). As an example, the mineralized region within the developing *tibia* of e16.5 embryos were significantly smaller and less mineralized compared to control embryos (Figure 1K). Upon closer examination, there were several developmental defects in the craniofacial region of *Cfdp1* conditional KO embryos, including hypo-mineralization of the skull, a smaller cranial vault, a reduced midface, and an increased cranial suture diameter (Figure 1M). Based on these observations, we conclude that *Cfdp1* is indispensable for mouse development and plays an essential role for gastrulation and early organogenesis in mice.

### CFDP1 associates with the mitotic spindle apparatus and is essential for physiological chromosome segregation

The embryonic lethality of *Cfdp1*^-/-^ mouse and previous studies demonstrating an essential role of CFDP1 in cell proliferation, ^[34,36,41]^ prompted us to examine CFDP1 function during mitosis in mammalian cells. Cell compartment fractionation studies followed by immunoblot analysis revealed high levels of CFDP1 in the chromatin-rich fraction and in the mitotic chromosome fraction of NIH3T3 cells (Figure S2A and S2B, Supporting Information). Immunofluorescence analysis using a monoclonal anti-CFDP1 antibody in NIH3T3 cells revealed a punctate staining pattern for CFDP1 within the interphase cell nucleus (Figure 2A). CFDP1 staining overlapped with DAPI dense foci in the nucleus in agreement with the role of CFDP1 in heterochromatin maintenance and function (Gopinathan et al, PLOS Biol 2024). We next visualized CFDP1 staining specifically in mitotic cells based on the high levels of CFDP1 observed in the mitotic chromosome fraction (Figure S2B). Immunofluorescence studies for CFDP1 in metaphase stage NIH3T3 cells revealed a striking staining pattern reminiscent of the microtubule network within the spindle assembly (Figure 2B). Furthermore, our double immunofluorescence staining experiments using CFDP1 antibody in conjunction with tubulin antibody confirmed that CFDP1 was associated with the mitotic spindle made of microtubules (Figure 2C). Interestingly, CFDP1 was also visualized at both centrosomes of the bipolar spindle assembly (Figure 2B, C). These immunofluorescence experiments indicated that CFDP1 associated with microtubules along the entire length of the spindle fibers from the centrosomes to the chromosomes aligned at the metaphase plate (Figure 2B, C).

**Figure 2.**
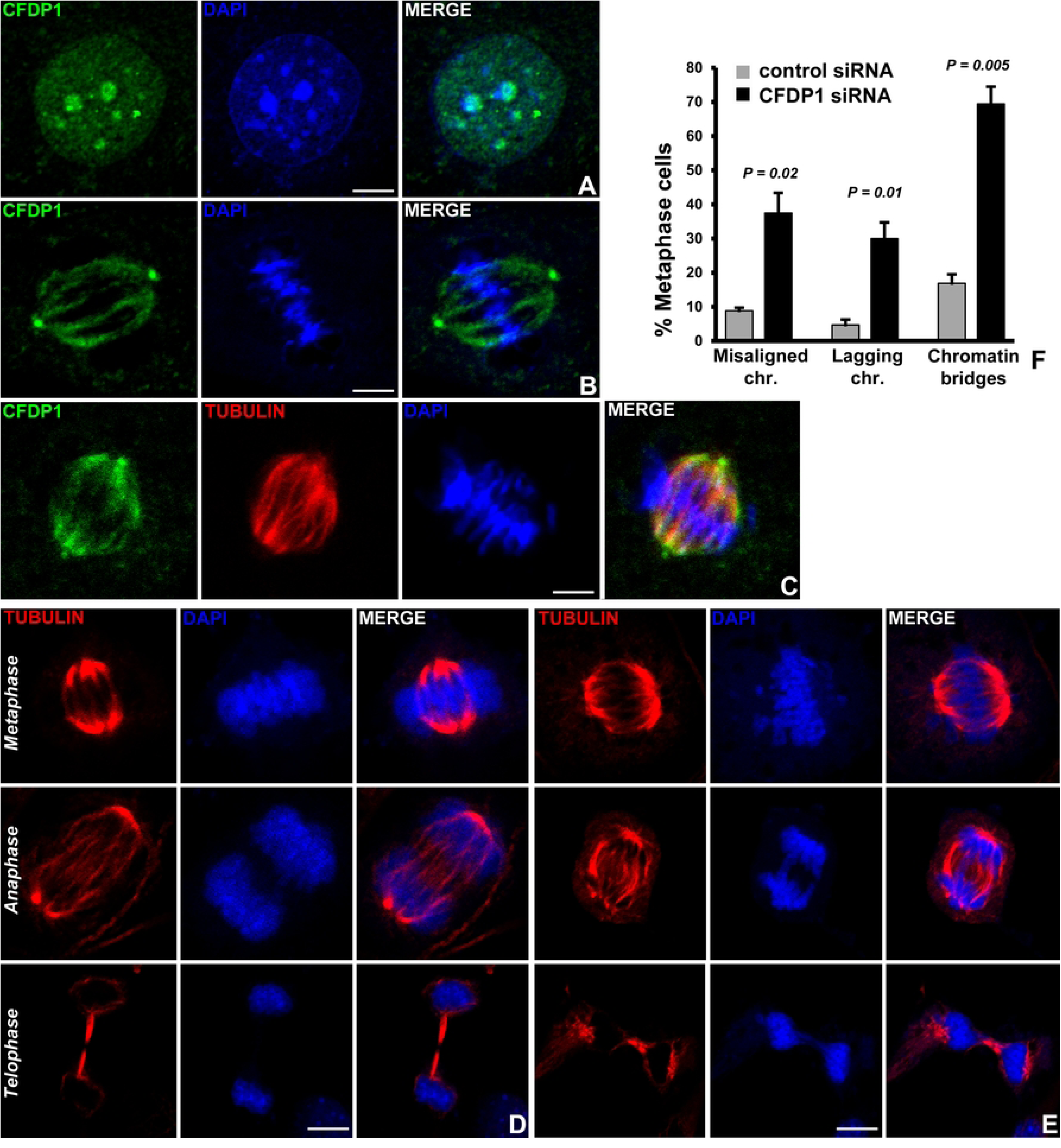
CFDP1 localizes to the mitotic spindle and CFDP1 deficiency leads to chromosome segregation defects. A) CFDP1 (green) is localized to the DAPI dense heterochromatin in interphase stage NIH3T3 cells. B-C) Immunofluorescence staining for CFDP1 localization in metaphase stage NIH3T3 cells using a monoclonal anti-CFDP1 antibody. CFDP1 staining can be visualized along the mitotic spindle and at the centrosome. Immunostaining was performed using CFDP1 antibody alone (B, green) or in combination with tubulin antibody (C, tubulin-red). Chromosome segregation defects in NIH3T3 cells treated with control siRNA (D) and CFDP1 siRNA (E). Various mitotic defects were visualized and scored after 72 hours of siRNA treatment. F) Quantitation of chromosome segregation defects in control and CFDP1 siRNA treated cells. A total of 200 metaphase stage, 120 anaphase stage and telophase stage mitotic cells were scored from three independent experiments (n=3). DNA is visualized with DAPI. Scale bars = 5 µm. *P* values were determined using unpaired student’s t test and considered significant when <0.05.

The association of CFDP1 with the spindle apparatus prompted us to ask whether CFDP1 was involved in chromosome segregation. CFDP1 protein level was downregulated in NIH3T3 cells using a pool of four specific small interfering RNA (siRNA SMART pool) directed against the coding sequence of mouse CFDP1 resulting in significant reduction of CFDP1 protein levels (Figure S2C, Supporting Information). We next performed immunofluorescence experiments to visualize the microtubule network of metaphase stage NIH3T3 cells treated with control or CFDP1 siRNA for 72 hours revealing striking defects in spindle organization in cells from the CFDP1 siRNA treated group (Figure 2D, E). Specifically, CFDP1 depleted mitotic cells exhibited a wide range of defects in the segregation of chromosomes which included failure of chromosomes to congress at the metaphase plate, high frequency of lagging chromosomes and anaphase chromosome bridges (Figure 2E) compared to control siRNA treated cells (Figure 2D). Interestingly, CFDP1 knockdown cells also displayed a high percentage of multi-pole spindles (Figure S2D, Supporting Information). The high incidence of lagging chromosomes in CFDP1 siRNA treated cells together with our immunofluorescence localization of CFDP1 at the mitotic spindle apparatus suggests an important role for CFDP1 in mitotic spindle organization and spindle stability.

### CFDP1 depletion affects K-fiber stability and triggers the spindle assembly checkpoint

The co-localization between CFDP1 and the mitotic spindle in conjunction with the chromosome segregation defects and abnormal spindle morphology after CFDP1 silencing prompted us to ask whether CFDP1 affects K-fibers as key stabilizing elements of mitotic spindles. Our studies demonstrated that K-fibers from CFDP1 depleted cells were shorter in length compared to controls, an effect that was observed both in cold stable, bipolar spindles (3.88 ± 0.93 µm in control *vs* 3.11 ± 0.98 µm in CFDP1 silenced cells, Figure 3A and 3B) and in cold stable monopolar spindles generated after Monastrol inhibition of the mitotic kinesin *Eg5* (3.07 ± 1.02 µm in control *vs* 2.35 ± 0.79 µm in CFDP1 silenced cells, Figure 3C and 3D). In addition, CFDP1 was found to be specifically localized to the minus end of the K-fibers in close proximity to the centrosomes of mitotic cells (Figure 3A).

**Figure 3.**
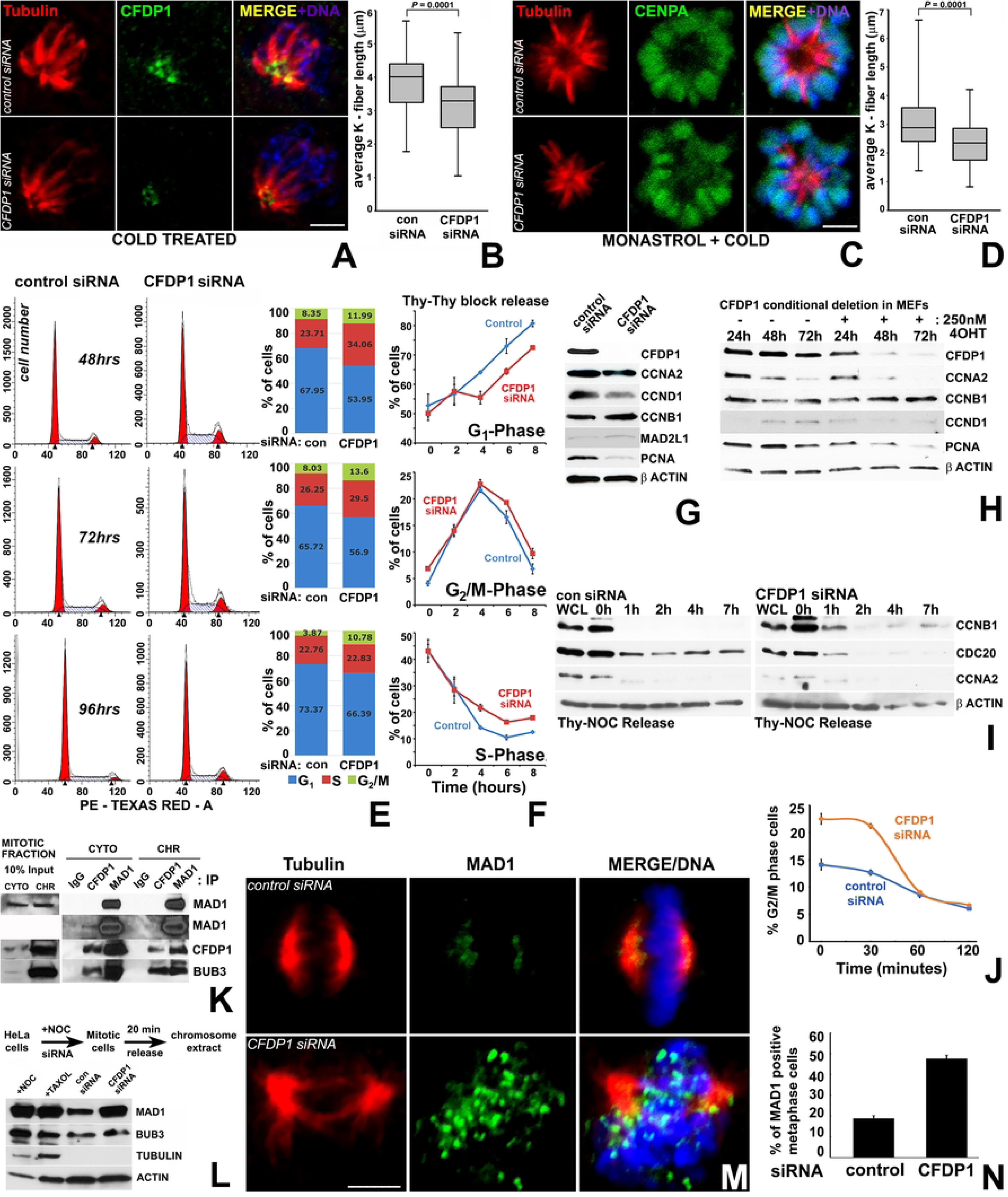
Defective K-fiber maturation and G2/M phase delay in CFDP1 knock down cells. A) K-fibers in cold treated control and CFDP1 knockdown NIH3T3 cells revealed by tubulin staining (red). CFDP1 is stained green. B) K-fibers were ∼20% shorter in CFDP1 depleted cells (data from n=150 cells). C) K-fiber instability in monopolar spindles of CFDP1 siRNA treated NIH3T3 cells after Monastrol and cold treatment. Tubulin and the centromere protein CENPA are stained red and green respectively. D) K-fibres were ∼23% shorter in CFDP1 depleted cells (data from n=150 cells). E, left) Cell cycle histograms after 48h, 72h and 96h of control and CFDP1 siRNA treatment. E, right) Quantitation of cells in G1, S and G2/M phases of cell cycle after control and CFDP1 siRNA treatment. Flow cytometry analysis was performed on cells from three independent experiments, and a representative profile is shown. F) Cell cycle progression in control and CFDP1 siRNA treated cells released from S phase block. Representative cell cycle progression is graphed (n=3). G) Cell cycle protein levels after 72h of control and CFDP1 siRNA treatment in NIH3T3 cells. H) Expression levels of cyclins and proliferation markers in 4-OHT (+) and vehicle treated (-, control) MEFs after 72h of induction of CFDP1 depletion as indicated. I) Delayed metaphase to anaphase transition upon CFDP1 depletion. Cyclin B1, A2 and CDC20 protein levels in control (left) and CFDP1 (right) siRNA treated cells after release from Thymidine-Nocodazole block. J) Flow cytometry analysis confirming delayed mitotic exit in CFDP1 depleted HeLa cells. Three independent experiments were conducted, and a representative profile is presented. K) CFDP1 immunoprecipitated spindle assembly checkpoint proteins, MAD1 and BUB3 from mitotic HeLa chromosomal lysates. CYT = cytoplasm, CHR = chromatin. L) Substantially higher protein levels of MAD1 but not of BUB3 in mitotic chromosome extracts from CFDP1 siRNA treated NIH3T3 cells compared to controls. M) Representative immunofluorescence analysis for MAD1 (green) on mitotic cells from CFDP1 depleted NIH3T3 cells. Tubulin is stained red and DNA revealed using DAPI (blue) staining. N) 30% increase in MAD1 positive metaphase stage cells upon CFDP1 depletion compared to control cels (n = 150 cells). Scale bars = 5 µm. Western blots from cell lysates and immunoprecipitation samples are representative of at least 3 independent experiments.

To determine whether chromosome segregation defects and abnormal spindle morphology in CFDP1 depleted cells impacts cell cycle progression, we performed flow cytometry analysis of asynchronously dividing NIH3T3 cells after CFDP1 siRNA revealing a higher percentage of cells in the G2/M phase and S phase of the cell cycle (Figure 3E). The effects of CFDP1 depletion on cell cycle progression were further verified in cells synchronized to early S phase using a double thymidine block. Upon release from S phase arrest, significantly more CFDP1 knock down cells were retained in the S and G2/M phase after 4, 6 and 8 hours (Figure 3F), indicative of a reduced number of cells completing the cell cycle after CFDP1 depletion. Consistent with the delay in cell cycle progression, expression of the cyclin D1 was reduced, while expression of the G_2_/M phase specific regulator, Cyclin B1^[42]^ was slightly elevated in lysates from CFDP1 siRNA treated cells (Figure 3G). Further supporting our concept of CFDP1 as an essential protein for cell proliferation, expression of the proliferation marker PCNA was greatly reduced upon CFDP1 knockdown (Figure 3G). To rule out potential side effects due to siRNA transfection, an alternative *Cfdp1* gene knockout strategy was carried out in tamoxifen (4-OHT)-treated inducible MEFs. This strategy was associated with a substantial decrease in CFDP1 expression after 72h of 4-OHT incubation (Figure 3H) and resulted in a drastic alteration of cell cycle modulators, including a decrease in Cyclin A2, Cyclin D1 and PCNA expression and an increase in Cyclin B1 levels after 48h and 72h (Figure 3H).

To identify the effect of CFDP1 on cell cycle modulators associated with M phase progression, control and CFDP1 siRNA treated NIH3T3 cells were released from an M phase block and analyzed for Cyclin B1, Cyclin A2, and CDC20 protein levels. Our analysis demonstrated elevated Cyclin B1 protein levels in CFDP1 siRNA treated cells over the time course analyzed (Figure 3I), suggestive of a decreased CyclinB1/Cdk1 complex degradation by the Anaphase promoting complex (APC),^[43]^ likely due to a prolonged M phase caused by a lack of CFDP1. Supportive of our interpretation, the APC activator CDC20 was elevated in control cells while its expression was reduced in CFDP1 knockdown cells at all time points analyzed (Figure 3I). Cyclin A2, which promotes G1/S and G2/M transition, was reduced in CFDP1 depleted cells when compared to control cells at the onset of our study and was no longer detectable at subsequent time points in either control or CFDP1 siRNA treated groups (Figure 3I). Furthermore, cell cycle analysis of siRNA treated NIH3T3 cells released from a metaphase arrest identified a significant percentage of cells corresponding to G2/M phase at early time points (0 and 30 min) compared to control cells, indicative of a prolonged M phase (Figure 3J). This delay in progression through M phase was due to cell-cycle check point activation, since our data demonstrated that CFDP1 interacted with two spindle assembly check point proteins, MAD1 and BUB3, in mitotic HeLa cells (Figure 3K).

Immunofluorescence analysis on CFDP1 siRNA treated NIH3T3 cells revealed extensive chromosomal targeting of the check point protein, MAD1 in metaphase cells (Figure 3M) and a 30% increase in MAD1 positive metaphase stage cells in CFDP1 siRNA cells compared to controls (Figure 3N). On a global level, the increase in chromosomal MAD1 levels upon CFDP1 depletion were confirmed by immunoblot analysis for MAD1 on chromosome extracts from mitotic cells (Figure 3L). Unlike MAD1, chromosome levels of BUB3 were not altered upon CFDP1 depletion (Figure 3L), suggesting that the role of CFDP1 in checkpoint activation might involve MAD1/MAD2 complex related mechanisms.

### The basic C-terminus and not the acidic N-terminus of CFDP1 protein interacted with tubulin, co-localized with the mitotic spindle, and promoted microtubule bundling and polymerization

Mammalian CFDP1 is a largely disordered protein comprising an acidic N-terminal region, a 40-aa lysine/glutamic acid/proline-rich repeat region and a highly conserved BCNT-C domain.^[44]^ The distinctive co-localization between CFDP1 and the microtubule network and its role in the regulation of K-fiber stability during mitosis suggested that CFDP1 might function as a Microtubule-Associated Protein (MAP). MAPs physically interact with tubulin and regulate microtubule dynamic instability for faithful chromosome segregation.^[45]^ We had previously demonstrated that only the acidic N-terminus region of CFDP1 interacted with histone H2A/H2B dimers *in vitro* (Gopinathan et al 2024). Comparing the N- and C-terminal regions of CFDP1 protein, exons 1 – 4 exhibited a low isoelectric point (pI < 5.0), while exons 5 – 7 were characterized by a relatively higher pI (> 8.0), indicating that CFDP1 has an acidic N-terminus and a basic C-terminus (Figure 4A). We hypothesized that CFDP1 polarity based on an extended isoelectric point (pI) range across the protein (2.8 for exon 1 and 9.0 for exon 7, Figure 4A) might explain its ability to interact with chromatin and microtubules (MTs).

**Figure 4.**
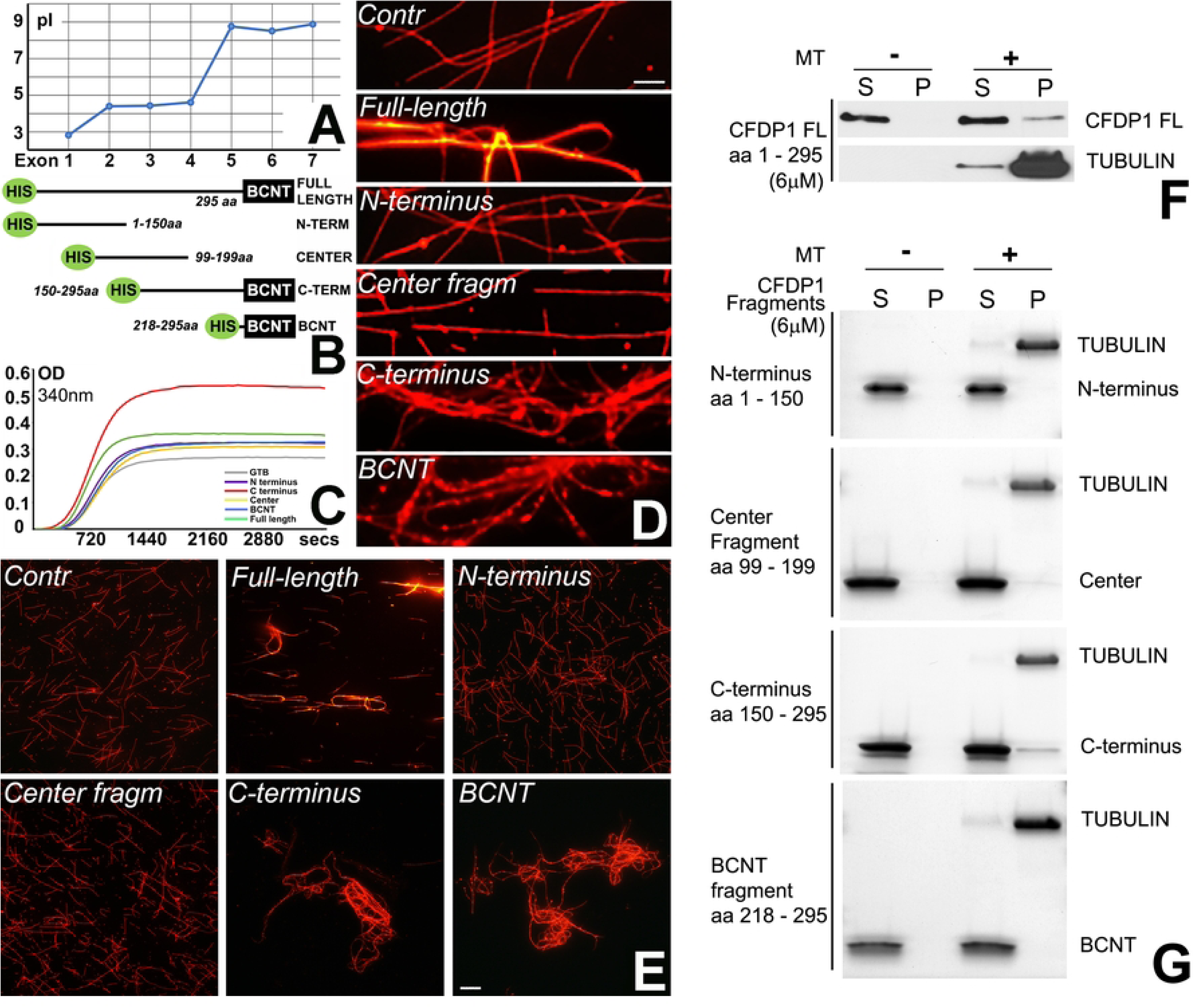
Microtubule bundling and polymerization are primarily mediated by the C-terminal basic half of CFDP1. A) Isoelectric point (pI) mapping of CFDP1 exons in mouse. B) Schematic representation of full length (FL) and truncated CFDP1 fragments used in the study. BCNT = BCNT domain. All fragments were produced with an N-terminal HIS tag (HIS). C) Tubulin polymerization assay with CFDP1 full length and CFDP1 fragments indicating robust tubulin polymerization by the C terminus of CFDP1. Representative kinetic graph is presented from five independent assays. D) High magnification micrographs and (E) overview of microtubule bundling observed in taxol-stabilized, rhodamine labeled microtubules incubated with full length and fragments of CFDP1. Images are representative of 5-6 independent MT bundling assays. F, G) Tubulin co-sedimentation assay with full length (F) and fragments of CFDP1 protein (G). Supernatant (S) and Pellet (P) fractions were run on an SDS-PAGE gel and stained with Biosafe Coomassie stain. The C terminus and to a lesser extent, the center fragment co-sediments with microtubules. Co-sedimentation assay is representative of two independent experiments.

To gain insights into the physiological effects exerted by the acidic and basic termini of CFDP1 on MT, four HIS tagged truncated fragments of the mouse CFDP1 were expressed: the acidic N-terminal fragment (amino acid (aa) 1-150), the basic C-terminal fragment (aa 150-295), the center fragment (aa 99-199) and the BCNT domain (aa 218-295) (Figure 4B). To determine if individual CFDP1 fragments affected tubulin polymerization dynamics, tubulin polymerization assays were carried out comparing the full length and individual CFDP1 fragments (Figure 4C). While the full length CFDP1 protein increased tubulin polymerization rate to a small extent, the C-terminal fragment resulted in a drastic increase in tubulin polymerization (Figure 4C, max OD at 340nm-control (GTB): 0.27; FL: 0.359; C-terminus: 0.541). Neither the N-terminal fragment, the center fragment nor the BCNT fragment exhibited noticeable levels of tubulin polymerization activity (Figure 4C). To test the ability of CFDP1 to self-assemble MTs into fiber bundles, either full length CFDP1 or its fragments were incubated with rhodamine labeled MTs. While incubation with the N-terminal fragment and the center fragment did not result in MT bundling, the full length CFDP1 protein resulted in modest tubulin bundling and both the CFDP1 C-terminus and the BCNT fragment exhibited robust MT bundling activity compared to controls (Figure 4D and 4E). We next investigated whether increased tubulin bundling and polymerization are due to a direct interaction between CFDP1 and MTs. We performed microtubule co-sedimentation experiments to detect the amount of CFDP1 full length or CFDP1 fragments that co-pellet with microtubules upon high-speed centrifugation. Appropriate controls were included for each CFDP1 test protein to detect any protein sedimentation in the absence of microtubules. Immunoblot analysis of the pellet fraction from full length CFDP1 incubated with microtubules demonstrated that CFDP1 specifically interacted with MTs and did not pellet in the absence of MTs (Figure 4F). Co-sedimentation experiments with CFDP1 fragments revealed that C-terminal fragment of CFDP1 co-sedimented with Taxol stabilized microtubules, indicating that MT-CFDP1 interaction was primarily mediated *via* the C terminal half of CFDP1. The center fragment also co-sedimented with MTs to a lesser extent, although it did not exhibit any bundling or polymerization activity in our previous assays (Figure 4F).

### CFDP1 interacted with TPX2 and was necessary for TPX2 mediated chromosomal microtubule nucleation

To further probe the molecular basis of CFDP1 function on the microtubule network, we conducted a Mass Spectrometric (MS) analysis of CFDP1 interacting proteins obtained by FLAG immunoprecipitation of lysates from NIH3T3 cells stably expressing 3xFLAG tagged CFDP1. Our assay identified members of the chromosome-dependent microtubule nucleation pathway, including TPX2, RCC1, RAN, Importins, tubulin proteins, kinesins and histone proteins as possible CFDP1 interacting proteins (Figure 5A and Table S2, S3, Supporting Information). The identification of RanGTP pathway members as CFDP1 interaction partners potentially explains the spindle defects observed in our CFDP1 depleted cells. To establish a role for CFDP1 in the chromosome-dependent MT nucleation pathway, we validated the interaction between CFDP1 and TPX2 in cytosolic and chromatin extracts from mitotic HeLa cells (Figure 5B) and between CFDP1, TPX2, tubulins α/γ and the nuclear protein import factors importins α/β in Hela mitotic extracts (Figure 5C). Since CFDP1 was specifically localized at the kinetochores of metaphase chromosomes, we also investigated whether CFDP1 interacts with members of the Chromosomal Passenger Complex (CPC), a kinetochore associated complex essential for MT stabilization and spindle assembly in mitotic cells.^[46]^ Our analysis revealed an interaction between CFDP1 and two members of the CPC, Aurora B kinase and INCENP (Figure 5C and Figure S3A, Supporting Information), further strengthening the link between CFDP1 and MT assembly. Interestingly, we did not detect an interaction between CFDP1 and the mitotic kinase, Aurora A or its active form - phospho Threonine 288 Aurora A (pT288 Aurora A), although both were robustly associated with TPX2 (Figure 5C). To precisely map the association between CFDP1 and TPX2 in mitotic cells, we performed immunofluorescence co-localization studies in mitotic NIH3T3 cells recovering from Nocodazole induced microtubule depolymerization as a means to distinguish centrosomal and acentrosomal microtubules.^[22]^ Our assays revealed that CFDP1 was exactly co-localized with TPX2 and tubulin at the microtubule nucleation sites on mitotic chromatin (Figure 5D, top panel and Figure 5E, left panel). CFDP1 exhibited a punctate staining pattern with intense staining at the microtubule nucleation sites 1 minute after nocodazole wash-out (Figure 5D top panel and Figure S4A, S4B, Supplemental Information) and stained more uniformly on the elongating microtubules at the 5-minute time point (Figure 5D, middle panel). In metaphase stage cells, CFDP1 was mostly concentrated at the centrosomes with diffused staining on the microtubule spindle apparatus (10-minute time point) (Figure 5D, bottom panel and Figure 5E, right panel).

**Figure 5.**
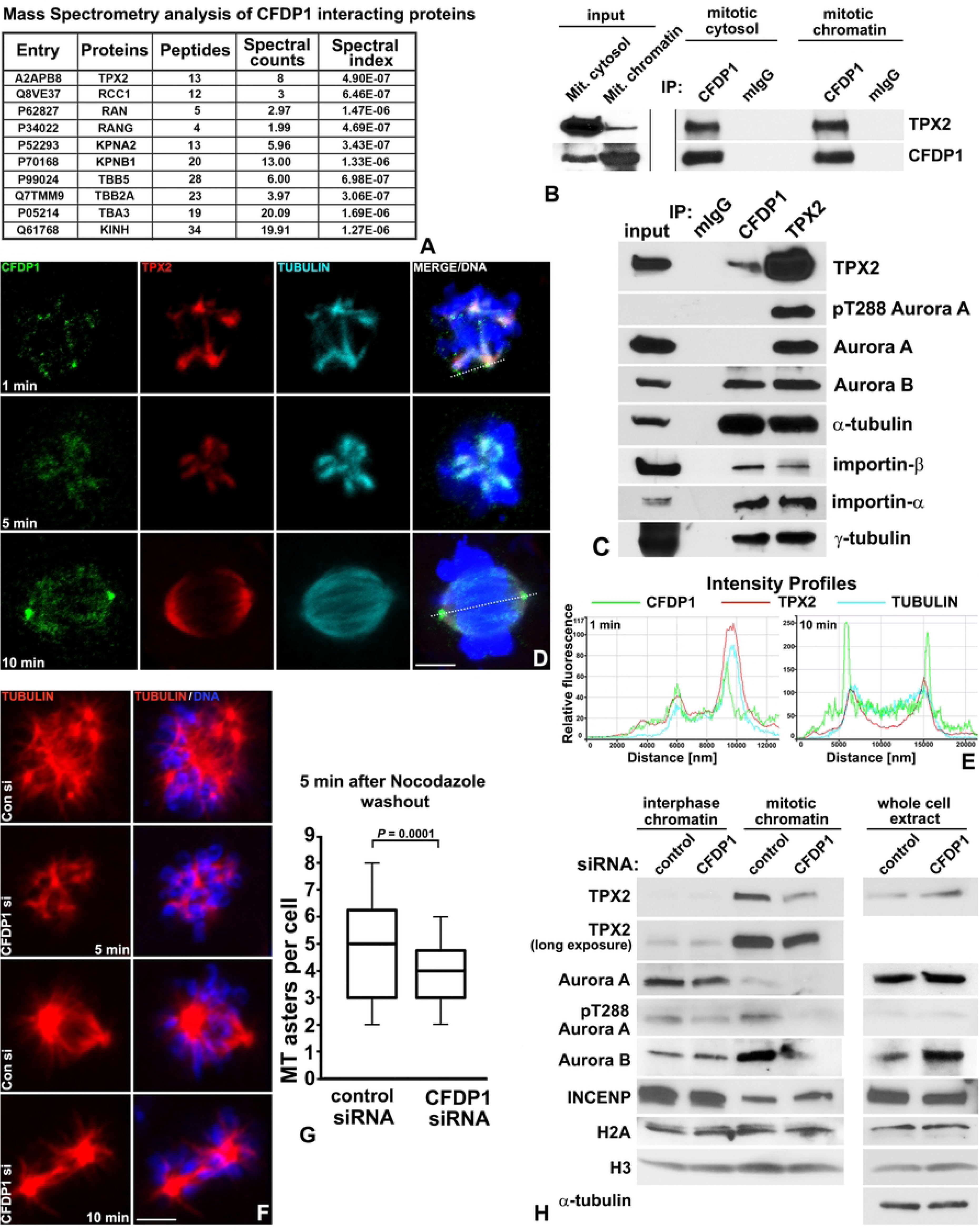
CFDP1 interacts with TPX2 and is essential for chromatin mediated microtubule nucleation. A) CFDP1 interacting proteins related to the chromosomal microtubule nucleation pathway identified by Mass Spectrometry analysis. B) CFDP1 interacts with TPX2 in cytosolic and chromatin extracts prepared from mitotic NIH3T3 cells. A control immunoprecipitation was performed using mouse IgG to determine the efficiency of pull-downs. C) Immunoprecipitation analysis of CFDP1 and TPX2 in mitotic HeLa cell extracts. CFDP1 interacted with TPX2 and α-tubulin but did not interact with Aurora A or its phosphorylated form (pT288 Aurora A). D) Representative immunofluorescence co-localization of CFDP1, TPX2 and tubulin in mitotic NIH3T3 cells released from nocodazole wash-out at the 1 min, 5 min, and 10 min time points. CFDP1 immunostaining (green) was closely associated with TPX2 (red) at the tubulin (blue) nucleation sites after 1 minute and at the later time point was localized to the centrosomes and tubulin. E) Changes in relative fluorescence signal intensity along the perforated line indicated in panel D for CFDP1 (green), TPX2 (red) and tubulin (blue) at 1 minute (left panel, nucleation phase) and 10 minutes (right panel, after spindle bi-orientation) after nocodazole washout. F) Representative immunofluorescence images visualizing microtubule (red) nucleation after nocodazole washout in control and CFDP1 siRNA treated NIH3T3 cells at the indicated time points. The Tubulin/DNA panel is a merge of both tubulin immunostaining with the corresponding DAPI staining for DNA. G) Number of MT asters per cell in control and CFDP1 siRNA treated cells 5 minutes after nocodazole washout. n = 80 cells for each condition. *P* valued was determined using unpaired student’s t test. H) Chromatin associated levels of select proteins from the microtubule nucleation pathway in control and CFDP1 siRNA treated NIH3T3 cells during interphase and mitosis (left panel). Total protein levels were assayed in whole cell extracts (right panel). Scale bars = 5 µm. Western blots are representative of 2-3 independent experiments.

Next, we tested whether CFDP1 is an essential component of the chromatin-dependent microtubule nucleation pathway by monitoring microtubule nucleation and changes in MT mass in control and CFDP1 siRNA treated cells. Five minutes after Nocodazole wash-out, control siRNA treated cells nucleated a large number of microtubule asters close to chromatin compared to CFDP1 knockdown cells (average of 4.83 ± 2.0 asters per cell in control vs 3.71 ± 1.3 asters in CFDP1 siRNA treated cells) (Figure 5F, 5 min panel and Figure 5G). After ten minutes, control cells organized a distinctive bipolar spindle, while CFDP1 knockdown cells lacked the typical organization of MT bundles radiating from each pole (Figure 5F, 10 min panel). We hypothesized that the decrease in chromatin mediated microtubule nucleation and MT mass in CFDP1 knockdown mitotic cells is a consequence of reduced TPX2 levels on the mitotic chromatin. To test this possibility, we analyzed levels of TPX2 in crude chromatin extracts prepared from control and CFDP1 siRNA treated interphase and mitotic cells. While chromatin levels of TPX2 were unaltered in interphase chromatin extracts, mitotic chromatin from CFDP1 knockdown cells exhibited a dramatic reduction in TPX2 levels (Figure 5H). The total levels of TPX2 were not affected upon CFDP1 siRNA treatment indicating that CFDP1 regulates TPX2 binding to chromatin. We also observed a significant decrease in chromatin levels of Aurora A, pT288 Aurora A and Aurora B in CFDP1 knockdown mitotic chromatin extracts implicating the RanGTP pathway as a potential target affected by the absence of CFDP1 (Figure 5H).

### CFDP1 promoted Aurora A activation and microtubule nucleation by decreasing importin α association with TPX2

The close association between CFDP1 and TPX2 at the microtubule nucleation sites prompted us to investigate whether CFDP1 and TPX2 co-bind microtubules in a complimentary fashion. Microtubule co-sedimentation assays revealed that consistent with our previous observations, CFDP1 co-sedimented with MTs while, addition of recombinant TPX2 protein did not significantly increase the binding of CFDP1 to MTs (Figure 6A, B). Surprisingly, at higher concentrations of TPX2, in spite of the increased TPX2 binding to MTs as observed in the pellet (P) fraction, CFDP1 binding to MTs was further reduced (Figure 6A, B). On the other hand, a gradual increase in CFDP1 protein level promoted TPX2 binding to MTs in co-sedimentation assays with a fixed concentration of TPX2 highlighting CFDP1 as a key regulator of TPX2-MT binding (Figure 6C, D). Since our previous assays have demonstrated both MT binding and bundling activities within the C-terminus region of CFDP1, we next investigated the binding between MTs and TPX2 in the presence of CFDP1 fragments. Co-sedimentation assays identified a robust increase in TPX2 association with MTs when incubated with the center fragment and the C-terminal fragment of CFDP1 (Figure S4A). Consistent with our previous observations, the N-terminal fragment and the BCNT domain fragment of CFDP1 were neither bound to MTs nor did they increase TPX2 binding to MTs (Figure S4A). We next confirmed the essential role of CFDP1 in MT assembly by performing MT aster assembly assays in HeLa mitotic lysates. While control lysates exhibited robust aster assembly leading to an increase in overall MT mass, lysates from CFDP1 immunodepleted extracts exhibited significantly less MT aster assembly and MT mass (Figure 6E). Importantly, MT aster assembly and MT mass could be reconstituted in CFDP1 immunodepleted mitotic lysates by the addition of exogenous CFDP1 protein in a dose dependent manner (Figure 6E).

**Figure 6.**
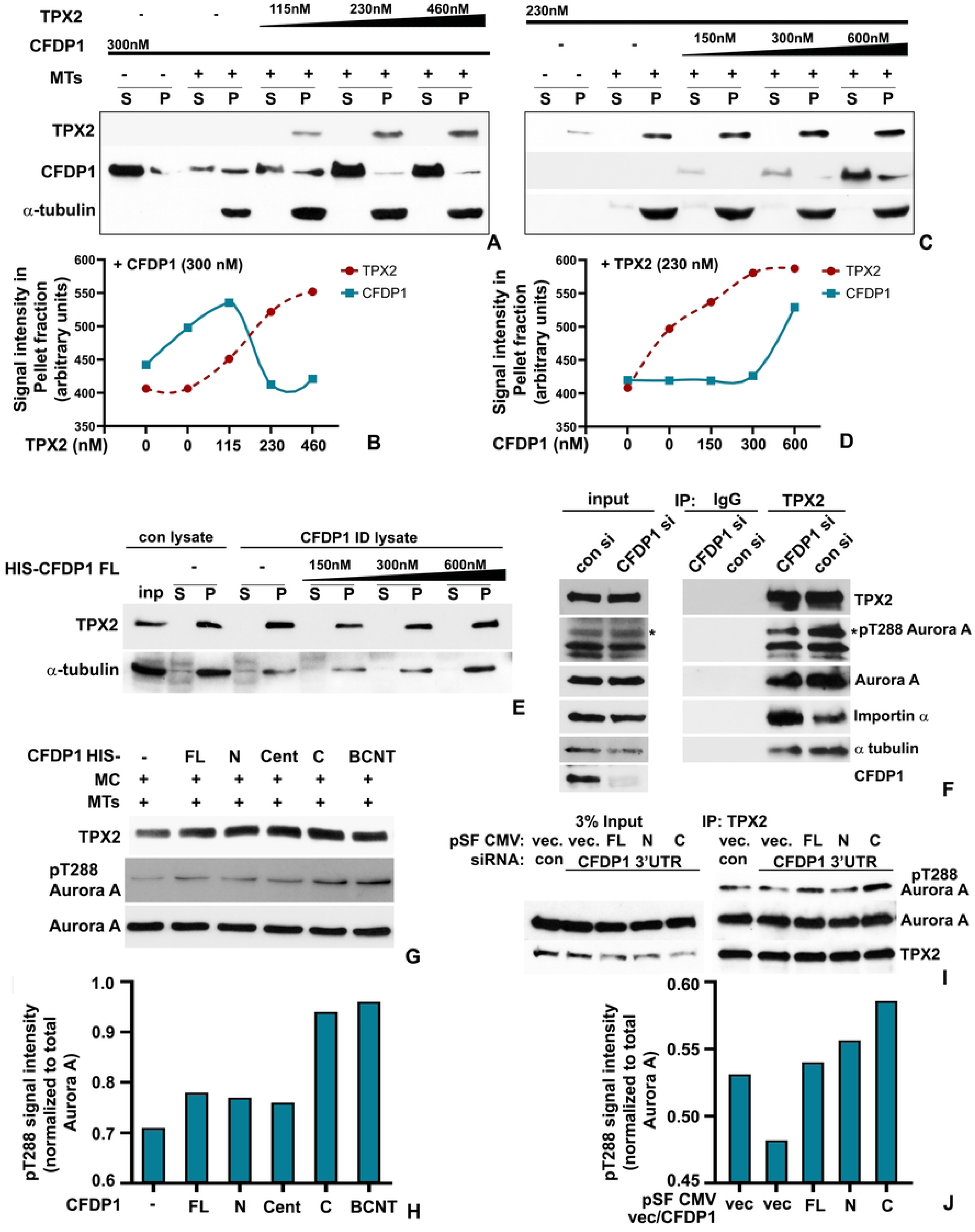
CFDP1 promotes microtubule (MT) assembly by activating Aurora A in a TPX2 dependent fashion. A-D) CFDP1 promotes TPX2 binding to microtubules. Addition of high concentrations of TPX2 protein decreased CFDP1 binding to MTs (A, B), whereas incubation with increasing concentrations of CFDP1 protein promoted TPX2 binding to MTs (C, D). MT co-sedimentation assays were performed with indicated concentration of proteins and revealed using western blot analysis. Representative results are shown from three independent experiments. E) CFDP1 is essential for *in vitro* microtubule assembly and addition of exogenous full length CFDP1 protein in CFDP1 immunodepleted mitotic extracts rescued microtubule assembly in a TPX2 dependent manner. Assay is representative of two independent experiments. TPX2 levels and MT mass were quantified by western blot assay. F) TPX2 from CFDP1 depleted cell extracts is associated with higher levels of importin α and lower levels of phosphorylated Aurora A (*pT288) and tubulin compared to control siRNA treated cell extracts. (n=3) G, H) The C-terminus and BCNT fragments of CFDP1 increases pT288 Aurora A levels *in vitro* in HeLa mitotic cytosol extracts. Cent = center fragment. Representative western blot derived from three independent experiments. I, J) Representative western blot demonstrating i*n vitro* rescue of Aurora A phosphorylation in HeLa cells co-transfected with siRNA directed against the 3’ UTR region of endogenous CFDP1 and plasmids expressing full length, N and C-terminus of CFDP1. (n=3). TPX2 immunoprecipitations from full length and C-terminus co-transfected mitotic lysates display rescue of pT288 Aurora A levels. S = supernatant, P = pellet, vec. = vector.

To determine whether TPX2 association with other members of the chromatin associated MT nucleation pathway was affected in the absence of CFDP1, we performed immunoprecipitation assays in mitotic lysates from CFDP1 knockdown cells. We used the well-documented interaction between TPX2 and Aurora A resulting in stabilization of Thr288-phosphorylation and activation of Aurora A as an indicator for MT nucleation and spindle formation efficiency.^[47]^ While, TPX2 from control siRNA treated mitotic lysates was stably associated with Aurora A and its phosphorylated active form pT288 Aurora A, in the absence of CFDP1, TPX2 immunoprecipitated significantly lower levels of both Aurora A and pT288 Aurora A (Figure 6F). Importantly, we found that in the absence of CFDP1, TPX2 was associated with significantly higher levels of the Ran-GTP pathway inhibitory protein, Importin α when compared to controls indicating that CFDP1 activates TPX2 by decreasing its association with Importin α (Figure 6F). Since our previous experiments identified the C terminus of CFDP1 as a key mediator of MT and TPX2 binding, we next investigated the physiological effects of this biochemical interaction between the C terminus, MT and TPX2 on Aurora A phosphorylation. Our *in vitro* Aurora A activation assays demonstrated that addition of the recombinant C terminal fragment of CFDP1 increased the levels of pT288 Aurora A in HeLa mitotic extracts (Figure 6G, H). Interestingly, in this assay, the BCNT fragment also exhibited a robust ability to increase pT288 Aurora A levels (Figure 6G, H). To further validate the role of CFDP1 C terminus as a key regulator of Aurora A activity, we assayed TPX2 mediated Aurora A phosphorylation in mitotic lysates prepared from HeLa cells following knockdown of endogenous CFDP1 mRNA (3’ UTR siRNA) and simultaneous re-expression of recombinant FL, N or C terminus fragments of CFDP1. Similar to our *in vitro* Aurora A activation assay, re-expression of the CFDP1 C terminal fragment in cells treated with 3’UTR siRNA for CFDP1 led to a dramatic increase in levels of TPX2 associated with pT288 Aurora A compared to control and CFDP1 siRNA treated cells (Figure 6I, J). In contrast, re-expression of N-terminal fragment of CFDP1 did not demonstrate any increase in levels of pT288 Aurora A associated with TPX2 (Figure 6I, J). Together, these rescue studies have confirmed the essential role of CFDP1 as a key regulator of TPX2 mediated microtubule nucleation and spindle formation.

### The acidic N-terminus of CFDP1 harbors a monopartite NLS essential for importin α dissociation from TPX2

Our previous experiments indicated a significantly higher level of TPX2 bound importin α in CFDP1 depleted cell lysates compared to controls. We therefore investigated whether CFDP1 directly activates TPX2 by destabilizing importin α-TPX2 interaction resulting in Aurora A phosphorylation and increased MT nucleation. Proteins harboring a high affinity classical Nuclear Localization Signal (cNLS) have been demonstrated to bind and sequester importin α leading to TPX2 displacement, freeing it to participate in spindle assembly.^[48]^ To determine whether CFDP1 functions through a similar mechanism, mouse CFDP1 protein sequence was scanned for presence of NLS using an NLS prediction software (cNLS Mapper). Our analysis revealed a putative monopartite NLS (amino acids 59-68) within the acidic N terminal of CFDP1 characterized by a short stretch of Arginine and Lysine rich basic amino acids (RKRK) (Figure 7A and Figure S4C, Supplemental Information). Similar basic amino acid stretches have been identified as a key requirement for NLS binding to importin α and for NLS function in several proteins.^[49]^ To test the function of CFDP1 NLS, we transfected GFP tagged N terminus, C terminus and FL CFDP1 into NIH3T3 cells and visualized them for nuclear translocation of the GFP tagged protein. As expected, GFP tagged FL CFDP1 (CFDP1-FL wt) clearly translocated exclusively to the nucleus while in cells expressing GFP vector alone, fluorescence was observed in both nucleus and cytoplasm (Figure 7B). On the other hand, while the acidic N terminus of CFDP1 comprising the NLS (CFDP1-N) translocated to the nucleus, mutating the RKRK motif to alanine (RKRK → AAAA, CFDP1-N mut) disrupted the nuclear import of the N terminal fragment with GFP fluorescence observed in both nucleus and cytoplasm (Figure 7A, 7B). Interestingly, both C terminus and CFDP1 FL with a mutated NLS (CFDP1-FL mut) were able to translocate to the nucleus effectively indicating that NLS activity as it related to nuclear translocation is not restricted to CFDP1 N terminus.

**Figure 7.**
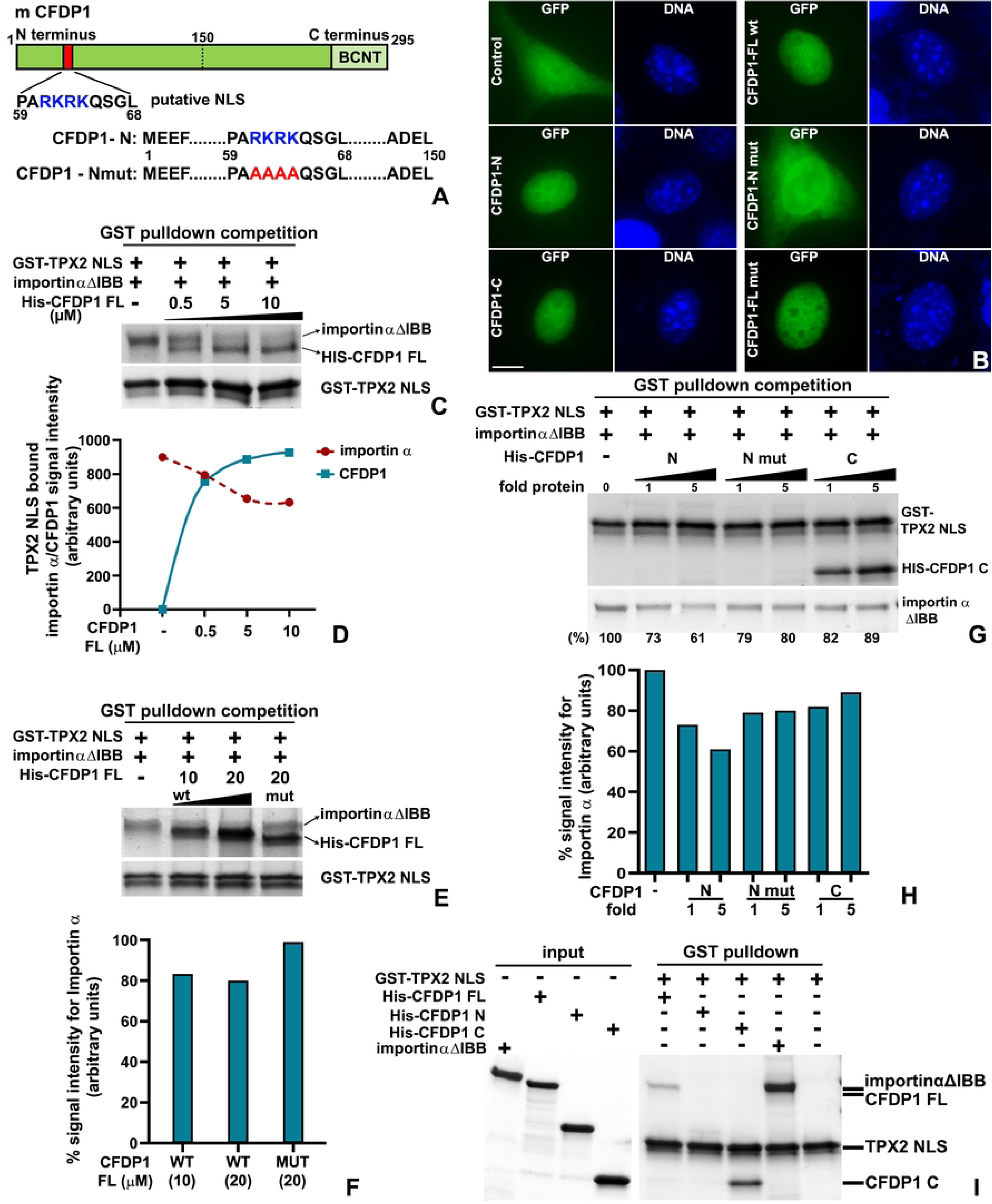
Direct interaction between CFDP1 and the TPX2 NLS promotes importin α dissociation. A) cNLS mapper analysis for putative NLS prediction in the mouse CFDP1 protein. NLS containing peptides are located at the N terminal end of CFDP1 (amino acid residues 59-68). Key NLS residues, RKRK (blue) within the putative NLS were mutated to AAAA (red) to generate the CFDP1-Nmut fragment. B) Representative images for GFP fluorescence (green) in NIH3T3 cells transfected with GFP tagged CFDP1 FL wt, N, N mut, C, FL mut constructs. Cells transfected with GPF vector (GFP) served as control. Scale bars = 5 µm. C, D) GST pulldown competition assay. Direct binding between CFDP1 FL and TPX2-NLS promoted importin α dissociation. Increasing molar concentrations of CFDP1 was added to preformed TPX2-NLS - importin α ΔIBB (Importin α lacking the IBB domain) complex and bound proteins were pulled down and analyzed by SDS-PAGE and Coomassie Blue staining. E, F) Comparison of importin α dissociation from TPX2 NLS in GST pulldown competition assays between CFDP1 FL wt and CFDP1 FL mut proteins. G, H) GST competition assay as in (C, E) with increasing concentrations of CFDP1 N, N mut and C terminal fragments as indicated. Addition of N terminal CFDP1 fragment resulted in a 39% reduction in TPX2 NLS associated importin α levels, compared to a 20% reduction with CFDP1 N mut. Both CFDP1 N and N mut fragments did not bind to the TPX2 NLS-importin α complex, while the C terminal fragment did (right two lanes, G). I) Demonstration of a direct interaction between the CFDP1 FL and the CFDP1 C terminus with the TPX2 NLS. The N terminal fragment exhibited negligible binding. All western blot-based assays were performed independently at least three times and representative blots are presented.

We next tested whether CFDP1 competes importin α off TPX2 by adding CFDP1 protein to preformed TPX2-importin α complex. For this analysis, we used the NLS containing domain of mouse TPX2 (TPX2-NLS, amino acids 302-322) which has been demonstrated to bind importin α Δ IBB (Importin α lacking the importin β binding domain, amino acids 75-496).^[48]^ *In vitro* GST pulldowns revealed that addition of increasing concentrations of full length CFDP1 to TPX2-importin α complex significantly reduced the levels of importin α bound to TPX2-NLS in a dose dependent fashion (Figure 7C, D). Additionally, concomitant with importin α dissociation in response to increasing CFDP1 concentration, CFDP1 exhibited progressive binding to the TPX2-NLS fragment (Figure 7C, D). In contrast, addition of an equimolar concentration of NLS mutated FL CFDP1 (CFDP1-FL mut) did not result in a similar decrease in TPX2 NLS associated importin α compared to CFDP1-FL wt (Figure 7E, F).

To specifically determine the contribution of N and C terminus of CFDP1 in destabilizing TPX2-importin α complex, we conducted GST pulldown competition assays with CFDP1 N, N mut and C terminal fragments. Our assays demonstrate that the addition of NLS containing N terminal fragment of CFDP1 decreased the levels of TPX2 bound importin α by 39% at the highest concentration tested (Figure 7G, H % values at bottom and quantitated in the graph) compared to only a 11% decrease in TPX2 bound importin α level with the addition of CFDP1 C terminus (Figure 7G, H). Importantly, the destabilizing effect of CFDP1 N terminus on TPX2-importin α complex was significantly lesser when CFDP1 N-mut fragments were used for GST competition with a 20% reduction in bound importin α levels (Figure 7G, H). Additionally, our assays demonstrate that consistent with our previous observation, the C terminal fragment of CFDP1 binds to TPX2-NLS, although, this binding did not result in any significant dissociation of importin α from the TPX2-importin α complex (Figure 7G). Furthermore, in support of this observation, we observed a direct interaction between CFDP1 FL and the C terminal fragment with TPX2-NLS while, the N terminal fragment exhibited almost negligible binding (Figure 7I). Together these results indicate unique and distinct roles for both N and C terminus of CFDP1 in NLS function, importin α dissociation from TPX2 and microtubule nucleation.

## Discussion

In the present study, we have used a broad range of approaches to probe the role of the CFDP1 chromatin protein as it relates to embryonic development and mitotic spindle formation. CFDP1 null mice were introduced to determine its role for embryonic survival. CFDP1 was localized to the mitotic spindle of dividing cells and siRNA mediated CFDP1 knockdown was performed to test the role of CFDP1 in cell cycle progression and K-fiber formation. Discrete CFDP1 N-terminal, center, C-terminal and BCNT fragments were generated to determine the effect of individual CFDP1 fragments on the structural and biophysical properties of microtubules. Co-immunoprecipitation and Mass Spectrometry assays were performed to examine the role of CFDP1 within the RanGTP pathway of microtubule nucleation. Focusing on TPX2 and Aurora A, interactions between CFDP1 and TPX2 triggering Aurora A activation were confirmed. Additional competition assays were performed to verify the role of the CFDP1 NLS binding region as it relates to TPX2 activation and MT nucleation. Together, our studies have established CFDP1 as an essential bipartite MAP that facilitates nuclear TPX2 activation through importin α dissociation and promotes chromosomal microtubule nucleation in conjunction with TPX2 and Aurora A.

Lethality of CFDP1 null embryos occurred remarkably early in mouse development. Based on the cell proliferation defects identified in CFDP1 knockdown studies we asked whether CFDP1 null phenotypes matched defects reported with the loss of other mitotic regulators. Confirming our lead, the timing of early embryonic lethality at e8.5 and cell proliferation defects in CFDP1 null embryos were strikingly similar to *Tpx2^-/-^* homozygous mutant mice.^[50]^ While Aurora A null embryos progressed until e10.5,^[51]^ they also exhibited spindle assembly defects similar to those observed in *Cfdp1* null embryos. *Cfdp1* null pre-implantation blastocysts resembled *Tpx2^-/-^* mutants and Aurora A null embryos in exhibiting growth arrest at the morula stage *in vitro*.^[50,51]^ Moreover, CFDP1 depletion in cells *via* siRNA led to abnormal spindle morphology, chromosome misalignment and a prolonged cell cycle. Together, the early embryonic lethality of *Cfdp1* null mice associated with defects in the spindle apparatus, chromosome segregation and cell cycle suggest that CFDP1 functions as a microtubule-associated protein (MAP) that results in similar loss-of function phenotypes as TPX2 and Aurora A.^[50,52,53]^

Our studies in mitotic cells detected CFDP1 on the mitotic spindle and centrosomes of metaphase stage cells and at the minus-ends of the K-fibers from cold-stable monopolar spindles. at three discrete locations: In metaphase stage cells, CFDP1 was strikingly associated with microtubules and the centrosomes indicating a key role for CFDP1 in organizing the mitotic spindle. However, CFDP1 association with microtubules along the entire length of the mitotic spindle was not detected in cold-stable K-fibers wherein CFDP1 was restricted to the K-fiber minus ends. The specific localization of CFDP1 at K-fiber minus ends and the shorter K-fibers in CFDP1 depleted cells support a concept of CFDP1 as a regulator of K-fiber maturation/stability. CFDP1 function may be similar to that of other K-fiber minus end microtubule associated proteins such as MCRS1, KANSL1, KANSL3 and the CAMSAPs/Patronin family proteins that play an important role in regulating K-fiber stability and are required for chromosome mediated microtubule assembly.^[54,55]^ Both MCRS1 and Patronin proteins regulate MT dynamics through the inhibition of MT minus-end disassembly by Kinesin-13 family of MT depolymerases. Suggestive of a role of CFDP1 in the inhibition of MT depolymerization at K-fiber minus ends, our mass spectrometry analysis identified two Kinesin-13 family protein members, Kif2a and Kif2c/MCAK as possible interaction partners for CFDP1. Through its dual localization at K-fiber minus ends and at the kinetochore, CFDP1 is ideally suited to not only stabilize K-fiber minus ends but also facilitate the formation of acentrosomal microtubules near the kinetochore, resulting in the promotion of K-fiber formation essential for poleward chromosome movement during mitosis. Since K-fiber mechanical integrity is critical for chromosome segregation, faulty K-fibers are the likely reason for erroneous chromosome segregation and M phase delay as a result of the improper kinetochore-MT attachment in CFDP1 depleted cells. In support of this concept, lysates from CFDP1 depleted cells revealed elevated levels of Cyclin B1 along with a concomitant decrease in CDC20 protein levels, indicative of a prolonged mitotic phase. Another consequence of the defects in microtubule nucleation and stabilization is the activation of the Spindle Assembly Checkpoint (SAC) following loss of CFDP1 since K fiber instability and unattached kinetochores have been reported to cause SAC activation.^[56]^

Our study demonstrated that the basic CFDP1 C-terminus and not the acidic N-terminus of the CFDP1 protein interacted with tubulin, promoted microtubule bundling and polymerization. There are two prominent regions within the CFDP1 C-terminus that may account for these robust microtubule bundling and polymerization properties, the lysine/glutamic acid/proline-rich 40 aa stretch intramolecular repeat (IR) region,^[44]^ and the highly conserved BCNT domain. Supportive of the CFDP1 IR region (aa 178-218) as a region involved in microtubule polymerization, the intrinsically disordered microtubule associated protein Tau contains a similar proline-rich region (PRR) within a C-terminal microtubule binding region (MTBR) involved in microtubule binding and polymerization.^[57]^ Thus, the CFDP1 IR domain might have similar roles related to the modulation of MT stability and dynamics as the Tau PRR/MTBR domain.

The second subdomain within the CFDP1 C-terminal region that may play a role in MT bundling and polymerization is the highly conserved BCNT domain. The BCNT domain was not only responsible for modulating microtubule structural properties in our MT bundling assays but also played a crucial role in chromatin binding in the Cfdp1 Drosophila homolog *Yeti*.^[37]^ A dual function involving both chromatin binding and MT bundling is not unique to the CFDP1 BCNT domain as such a dual role has also been attributed to several other MAP-type chromatin proteins.^[32,33]^ Increased microtubule bundling and polymerization rates facilitate mitotic spindle formation,^[24,25,58,59]^ suggesting that CFDP1 functions as a chromatin MAP enabling mitotic chromatin spindle formation through its basic C-terminus. Microtubule bundling and stability are essential for K-fiber assembly, chromosome segregation and cell division,^[54]^ and loss thereof as it occurs in CFDP1 depleted cells explains the spindle and K-fiber defects and mitosis phenotypes observed in our CFDP1 depleted cells and embryos. The acidic N terminal fragment of CFDP1 did not exhibit any interaction with microtubules in our assays and was shown to interact with histone H2A-H2B *in vitro*. In support of a chromatin-related function of the CFDP1 N-terminus, the acidic N terminal domain of the CFDP1 yeast homologue Swc5 preferentially binds to the histone H2A-H2B dimer facilitating SWR mediated H2A.Z variant exchange within the yeast nucleosome.^[60]^

Our study demonstrated that CFDP1 is necessary for TPX2 mediated chromosomal microtubule nucleation by decreasing importin α levels association with TPX2 and promoting Aurora A activation. Chromosomal microtubule nucleation is an essential process that allows for mitotic spindle assembly in addition to the centrosomal spindle assembly pathway. The chromosomal microtubule nucleation pathway relies on a RanGTP gradient generated by chromosome-bound RCC1 that supports activation of spindle assembly factors such as TPX2 by promoting its dissociation from the karyopherin importin α. TPX2 in turn activates the Aurora A kinase, which then phosphorylates the adapter protein NEDD1 on Ser405 within the XRHAMM-NEDD1-γ-TuRC complex, fulfilling an essential step for Ran-GTP dependent MT nucleation.^[61,62]^

A nocodazole washout revealed CFDP1 together with TPX2 and tubulin within the central core region of emerging microtubule asters, providing further evidence for CFDP1 as a facilitator of chromosome-driven MT nucleation. Our experiments documenting promotion of MT polymerization by CFDP1 and an overall decrease in the number of MT asters upon CFDP1 depletion argue for a role for CFDP1 during both the initial seed stage and the subsequent elongation stage of MT nucleation in the aster. In support of an early function in MT nucleation, CFDP1 interacted with gamma-tubulin, an indispensable component of MT-organizing centers (MTOCs) that regulate *in vivo* MT nucleation and organization in all eukaryotes.^[63]^ The most likely explanation for the effect of CFDP1 on MT nucleation and formation is its highly disordered protein structure,^[44]^ which allows it to form liquid-like drops through a demixing process reported for several other intrinsically disordered proteins such as the neuronal MAP Tau. These polyproline-rich domain containing proteins then cause phase separation and formation of tau drops inside which microtubules are nucleated as a result of macromolecular crowding and increased local tubulin concentration. ^[64,65]^

TPX2 has long been known as a crucial regulator of local MT nucleation in close proximity to the mitotic chromosomes. ^[52,66,67]^ However, it is not clear how TPX2 interfaces with mitotic chromatin on a physical level. Here we introduce CFDP1 as a TPX2 MT nucleation partner as it enriches TPX2 on the mitotic chromatin. We propose that this physical tethering of TPX2 to the mitotic chromosomes as a result of its interaction with kinetochore bound CFDP1 sets the stage for non-centrosomal MT formation at the kinetochore where MTs are rapidly bundled to form K-fibers. Such a scenario has been supported by several *in situ* studies demonstrating that short and randomly oriented non-centrosomal MTs occur in the immediate vicinity of the centromere, leading up to end-on attachment with the kinetochore.^[19,21,22,68]^ Based on these studies and our data, CFDP1 emerges as a MAP functioning in multiple steps of MT nucleation and assembly near chromosomes, beginning with the stabilization of an initial MT seed and thereafter facilitating rapid MT elongation in a TPX2 dependent fashion.

Our data demonstrate that CFDP1 regulates at least two aspects of TPX2-mediated MT nucleation and assembly during mitosis. First, we identified that CFDP1 promoted TPX2 binding to MTs in a dose dependent fashion. Electron microscopy studies have revealed that TPX2 employs two flexible MT-interacting elements (ridge and wedge) to bind tubulin in a critical step to initiate MT nucleation.^[69]^ *In vitro* experiments have also demonstrated that TPX2 directly promotes MT stability by reducing the frequency of catastrophes and/or by stabilizing MT nucleation intermediates.^[70,71]^ Given the MT related functions of CFDP1 and its close association with TPX2 at the microtubule nucleation site, CFDP1 may contribute to either of these steps, resulting in the stabilization of MT-TPX2 interactions and MT nucleation intermediates. One interesting aspect of TPX2 interaction with microtubules is its higher affinity to MT ends with a characteristic curvature.^[71]^ In our bundling assays, CFDP1 affected MT structure to form such curvatures, suggesting that CFDP1 modulates MT physical properties to enhance TPX2 binding at nucleation sites.

The second aspect by which CFDP1 functions during TPX2-mediated MT nucleation and assembly is the conserved TPX2-Aurora A pathway, regulating Aurora Kinase A activation by TPX2. Release of TPX2 from importin-α/β in the proximity of chromosomes in a RanGTP mediated fashion allows TPX2 to interact with and activate Aurora A.^[22,47,62,72]^ Our experiments suggest that CFDP1 plays a crucial role in the early step of the MT nucleation cascade by counteracting importin α-mediated TPX2 sequestration, resulting in Aurora A activation. Lysates from CFDP1 siRNA treated cells yielded significantly lower levels of TPX2-associated phosphorylated Aurora A (pT288 Aurora A), explaining the MT nucleation defects and spindle abnormalities observed in cells lacking CFDP1. It is not clear as of right now whether CFDP1 acts in parallel or synergistically with RanGTP to activate TPX2 and induce MTs near mitotic chromatin. Our study suggests that CFDP1-mediated TPX2 activation may serve as an alternate mechanism driving microtubule assembly adjacent to chromosomes in addition to the RanGTP pathway. Regardless of whether CFDP1 directly activates TPX2-mediated MT nucleation or does so in conjunction with RanGTP, both functions related to TPX2-MT interaction and tubulin structure modulation are mediated by the CFDP1 C-terminus, lending further support to the concept that the CFDP1 C-terminus acts as the primary modulator of MT nucleation.

The acidic N terminus of CFDP1 harbors a monopartite NLS essential for importin α dissociation from TPX2. Sequestration of importin α by highly concentrated and localized nuclear localization signal (NLS)-containing proteins during mitosis has been suggested as a general mechanism for the activation of Spindle Assembly Factors (SAFs) such as TPX2.^[48]^ However, regardless of the biochemical interaction between CFDP1 and importin α/β in mitotic lysates, we did not detect a direct interaction between CFDP1 and importin α *in vitro* (data not shown), suggesting that the CFDP1 N terminus harboring the NLS dissociates importin α from TPX2 *via* a non-competitive mechanism. Moreover, the addition of increasing amounts of the N-terminal fragment to TPX-2 NLS bound importin α resulted in increased dissociation of importin α from TPX2, suggested a concentration-dependent mechanism. According to recent studies, TPX2 is comprised of a total of 3 NLS regions potentially mediating importin α interactions (NLS1, NLS2 and NLS3).^[73]^ So far, we have only tested the TPX2 NLS2 fragment in our competition assay, and NLS1 and NLS3 might interact differently with the CFDP1 N-terminus, affecting importin α binding to TPX2. We speculate that CFDP1-mediated preferential control of TPX2 activation through individual TPX2-NLS regions (NLS1 vs. NLS2 vs. NLs3) might fine tune TPX2-mediated MT nucleation during mitosis, allowing it to seamlessly integrate into the RanGTP pathway.

Together, these studies suggest that CFDP1-mediated TPX2 activation and MT nucleation/elongation is a multistep process (Figure 8). Based on the substantial differences between CFDP1 C-terminal and N-terminal TPX2-NLS binding, we propose that initially, CFDP1 recruits the TPX2/importin α complex at the kinetochore region of mitotic chromosomes. As a second step, the N-terminal region of the TPX2-bound CFDP1 dissociates importin α from the TPX2-NLS, freeing TPX2 to initiate MT nucleation (as illustrated in Figure 8B and 8C). In the next step, after activating TPX2, CFDP1 exerts its microtubule binding and bundling activities *via* its C terminus to facilitate MT assembly near the kinetochore in a TPX2 dependent fashion (as illustrated in Figure 8E and 8F). Such a step-wise MT nucleation and elongation procedure would focus the TPX2 mediated MT nucleation process on the immediate proximity of the kinetochore. By virtue of its location at the kinetochore, CFDP1 is ideally situated to promote MT nucleation in the immediate vicinity of the kinetochore region where microtubules are easily captured and stabilized by the kinetochore complex to form K-fibers. K-fiber formation favors a “search and capture” mechanism of spindle formation that supports chromosome attachment and biorientation ^[74]^, possibly mediated by CFDP1.

**Figure 8.**
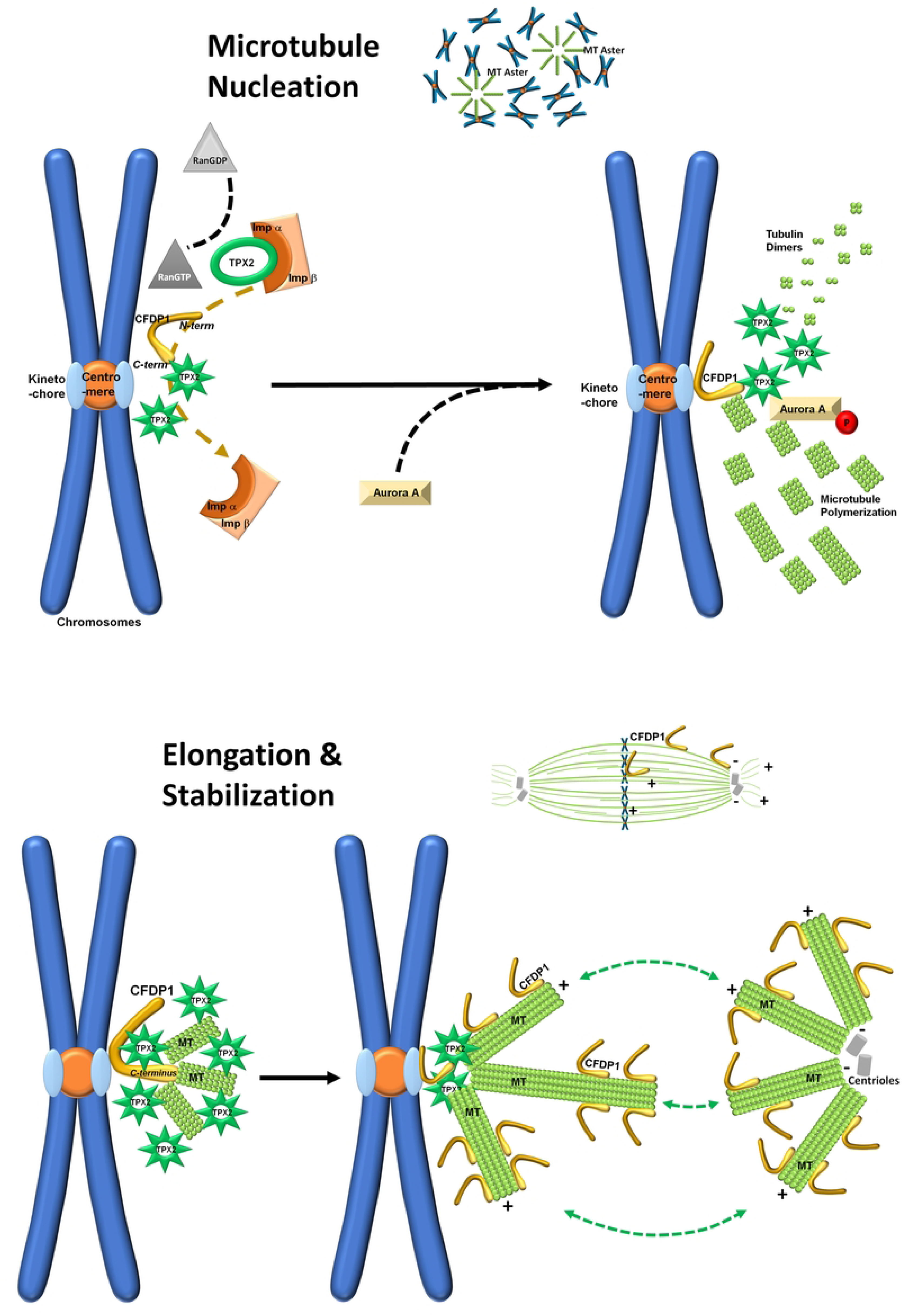
Sketch illustrating CFDP1-mediated acentrosomal MT assembly in mitosis. A) Acentrosomal MT nucleation and aster formation (MT Aster) in the immediate vicinity of mitotic chromosomes. B) The N-terminus of kinetochore-bound CFDP1 promotes local dissociation of TPX2 from its inhibitory interaction with importin α/β. C) Following its release and activation, TPX2 in turn activates Aurora A, resulting in a stimulation of MT nucleation facilitated by the CFDP1 C terminus near the kinetochore. D) CFDP1 localization in relationship to the spindle apparatus and the metaphase plate during metaphase. E) MTs nucleated near the kinetochores are captured by CFDP1 and TPX2. F) CFDP1 associates with and stabilizes the spindle MTs leading to K fiber formation (double arrows).

## Materials and Methods

### Generation of *Cfdp1* knockout mice

All animal procedures were approved by the Institutional Animal Care and Use Committee of the Texas A&M University Health Science Center. *Cfdp1* knockout mice (*Cfdp1*^-/-^; referred as *Cfdp1* KO) and *Cfdp1* conditional knockout mice (*Cfdp1*^flox/flox^; referred as *Cfdp1* conditional KO) were generated independently in the C57BL/6 (B6) genetic background using the targeting strategy described in Supplementary materials. For the generation of *Cfdp1* inducible knockout mice, B6.129 *Gt(ROSA)26Sor^tm1(Cre/ERT2)Tyj^*/J (JAX 8463) homozygous mice (Jackson Laboratories) were crossed with mice heterozygous for *Cfdp1* KO allele (*LacZ - Neo* insertion). Males from the resulting F1 progeny (*ROSA26-Cre*^+/-^; *Cfdp1*^+/-^) were crossed with female *Cfdp1*^flox/flox^ mice to generate test mice (*ROSA26-Cre*^+/-^; *Cfdp1* exon1^flox/-^) and control mice. Conditional deletion of *Cfdp1* was carried out *in utero* in mouse embryos by administering 2 mg Tamoxifen (Sigma, dissolved in corn oil) i.p. to pregnant mice (at e8.5 of gestation) and embryos harvested after 4 days for genotyping and phenotypic analysis.All studies were performed either in male or female mice and no gender specific phenotypic differences were observed. Genotyping for identification of *Cfdp1* alleles was performed with specific oligonucleotides (Table S1) as described in Supplementary materials. Embryos were staged based on the first day of vaginal plug, denoted e0.5.

### Cell lines and Primary cell cultures

NIH3T3 cells (CRL-1658) and HeLa cells (CCL-2) were obtained from the American Type Culture Collection (ATCC) and maintained in DMEM (Sigma) high glucose media with 10% FBS at 37°C in a humidified atmosphere of 5% CO_2_. Mouse Embryonic Fibroblasts (MEFs) were isolated from e13.5 embryos. Briefly, head and visceral organs were removed and remaining embryonic tissue minced finely with a scalpel blade. Tissue pieces were digested with 1X Trypsin EDTA (GIBCO) for 30 min at 37°C, and cells plated in DMEM supplemented with 10% FBS and 1X antibiotics. *Cfdp1* conditional deletion in MEFs was initiated by adding 250 nM 4-hydroxy Tamoxifen (4-OHT, Millipore; dissolved in 100% Ethanol) to culture media and cells analyzed at indicated time points.

### Histology

WT and *Cfdp1* KO embryos were fixed in 10% neutral buffered formalin and embedded in paraffin wax. 6 µm sections were cut and stained with hematoxylin-eosin or subjected to immunohistochemistry using Histostain plus Broad Spectrum kit (Life Technologies). Sections were incubated with primary antibodies overnight at 4°C, treated with HRP conjugated secondary antibodies and protein localization revealed using the AEC Red kit (Life Technologies). For cell proliferation assays, BrdU (Sigma) was injected (100 mg/kg body weight) in pregnant mice on the 6^th^ gestational day. Embryos were harvested after 2 hours, fixed in Carnoy’s fixative, dehydrated and embedded in paraffin wax. BrdU incorporation was revealed using BrdU IHC Kit (EMD Millipore).

### Cell synchronization

S phase and M phase synchronization were carried out by a double Thymidine block or Thymidine-Nocodazole block respectively. For double Thymidine block, exponentially growing cells were incubated with 2.5 mM Thymidine (Sigma) for 16 h. After Thymidine wash off, cells were allowed to recover for 10 h followed by a second Thymidine block (2.5 mM) for 16 h. Cells were subsequently harvested for flow cytometry. M phase synchronizations were performed by Nocodazole (Sigma, 100 ng/ml for 12-16 h) or by a Thymidine-Nocodazole block (Nocodazole added after the first Thymidine block). Mitotic cells were harvested by manually shaking the dishes. Synchronized cells were released from respective blocks and harvested for flow cytometry analysis.

### Flow Cytometry sample preparation and data acquisition

Control and experimental cells were trypsinized with 1X Trypsin EDTA, washed with 1X PBS and ∼ 100,000 cells were fixed in cold −20°C, 70% Ethanol and stored at −20°C until staining. For data acquisition, fixed cells were washed with 1X PBS and suspended in 1X PBS containing 0.5% BSA, 40 µg/ml Propidium Iodide (Sigma) and 40 µg/ml RNase A (ThermoFisher Scientific). Samples were incubated at 37°C for 15 min and data acquired on a BD FACSCalibur flow cytometer (BD Biosciences) at an event rate of < 500 events/second (Supplementary materials).

### Construction of plasmids

Coding sequence for full length mouse *Cfdp1* (885bp) and fragments, N-terminus (1-450 bp), C-terminus (451-885 bp), Center fragment (297-597 bp) and the BCNT fragment (654-885 bp) were synthesized with a 5’ 6X HIS tag using Platinum *Taq* DNA Polymerase (ThermoFisher Scientific) from NIH3T3 cDNA and cloned in pEXPR-IBA105 (IBA) for mammalian expression and in pASK-IBA43plus (IBA) for bacterial expression. For the Mass Spectrometry study and rescue study in NIH3T3 cells, *Cfdp1* was cloned in psF-CMV-NEO-NH2-3XFLAG (Sigma) resulting in an N terminal 3X FLAG fusion protein. The NLS mutated N-terminus fragment of CFDP1 was generated by site directed mutagenesis using the Q5 site-directed mutagenesis kit (NEB) and specific primers. For Nuclear localization studies, FL, FLmut, N, Nmut and C terminal fragments of CFDP1 were cloned in pAcFFP1-C1. Mouse TPX2 NLS sequence (nucleotides: 903-966) cloned in pGEX-2TK with a N-terminal GST tag was obtained from Origene. Oligonucleotides are listed in Table S1.

### Expression and purification of proteins

pASK-IBA43plus vectors coding HIS tagged *Cfdp1* constructs and pGEX-2TK vector coding TPX2 NLS were transformed into chemically competent *E. coli* BL21(DE3) (ThermoFisher Scientific) by a heat-shock process. Bacterial cells were selected overnight at 37°C on Luria Bertani (LB)-agar plates supplemented with 100 µg/ml Ampicillin. A single resistant colony was cultured overnight at 37°C in LB/Ampicillin. Pre-cultures were diluted 100-fold in 25 ml of LB/Ampicillin and grown for 3 h at 37°C until an absorbance of 0.5 at 600 nm was attained. Protein expression was induced with Anhydrotetracycline (IBA) (200 µg per liter) for HIS tagged proteins and 0.1 mM IPTG for GST tagged proteins and further grown for an additional 3 h at 37°C. Cells were subsequently collected at 5000 g for 10 min in 5ml aliquots and stored at −80°C. HIS tagged proteins were purified under native conditions using Ni-NTA Spin Kit (Qiagen) as per manufacturer’s instructions. Bacterial pellet from a 5 ml induced culture volume was processed for each spin column purification. GST tagged protein was purified using the MagneGST Protein Purification System (Promega) following manufacturer’s instructions. Desalting and buffer exchange (80 mM Hepes, pH 8.0, 1 mM EGTA, 1 mM MgCl_2_, 1 mM DTT) was carried out in Amicon Ultra centrifugal filters (Millipore) with a 3 kDa cutoff membrane. Protein concentration and integrity was verified by SDS-PAGE and staining with Colloidal Coomassie stain (Bio-Rad). Full length human TPX2 protein (TP305821) and mouse importin α Δ IBB (70-529 aa, untagged) were obtained from Origene and used in binding assays after buffer exchange.

### Mammalian cell transfections and siRNA treatments

Transfections were performed using Lipofectamine 3000 (ThermoFisher Scientific) and transfected cells selected using G418 (Gibco). Short interfering RNA oligonucleotides (SMARTpool siRNA, Horizon Discovery) targeting mouse and human CFDP1, and scrambled siRNA (control siRNA) were introduced into cells at a concentration of 75 nM using DharmaFECT 1 reagent (Horizon Discovery) following manufacturer’s instructions. For siRNA and plasmid co-transfection experiments, cells were treated with a mixture of siRNA (targeting the 3’ UTR of mouse or human CFDP1) and plasmid constructs expressing CFDP1 using the DharmaFECT Duo transfection reagent (Horizon Discovery).

### Tubulin assays

For Tubulin polymerization assays, 100 µl of a 3 mg/ml Tubulin stock (Cytoskeleton) prepared in General Tubulin Buffer (GTB; 80 mM PIPES pH7.0, 2 mM MgCl_2_, 0.5 mM EGTA) containing 1 mM GTP and 10.2% glycerol was incubated with CFDP1 full length or fragment proteins (diluted to 10 µM in GTB) at 37°C and absorbance measured at 340 nm using a spectrophotometer (SpectraMax 250) set in kinetic absorbance mode. Measurements were acquired once every minute for a total of 60 min.

For microtubule (MT) bundling assays, MTs were assembled from TAMRA Rhodamine labeled Tubulin (Cytoskeleton) in GTB with 10% Glycerol and 1 mM GTP at 37°C for 20 min. MTs (4 mg/ml) were further diluted 1:200 in GTB containing 20 µM Taxol (Cytoskeleton) and 6 µl of it incubated with 5 µM of CFDP1 full length or fragment proteins in a 10 µl reaction at room temperature for 20 min. Reaction mixtures were placed under a coverslip immediately and imaged under a fluorescence microscope equipped with a 585 nm emission filter (Leica DMRX).

For MT co-sedimentation assays, MT assembly was carried out in GTB as above in the presence of 1 mM GTP at 35°C for 20 min. Polymerized MTs were stabilized using Taxol (20 µM) and diluted in GTB containing Taxol to a concentration of 5 µM and stored at room temperature. Test proteins (5 µM) were incubated with 20 µl of Taxol stabilized MTs for 30 min at room temperature. Reaction mixtures were centrifuged at 100,000 g for 40 min at 25°C through a Glycerol Cushion Buffer (60% Glycerol in GTB) and supernatant and pellet fractions analyzed using SDS-PAGE. Specificity of MT sedimentation assays was monitored by analyzing the pellet fraction after incubating CFDP1 or TPX2 proteins in the absence of microtubules.

### *In vitro* microtubule aster assembly

Mitotic HeLa cells collected by shake-off were incubated with 20 µg/ml cytochalasin B for 30 min at 37°C. Cells were washed twice with cold 1X PBS and once with cold KHM buffer (78 mM KCl, 50 mM Hepes pH 7.0, 4 mM MgCl2, 2 mM EGTA, 1 mM DTT, 1X Protease inhibitors) in the presence of cytochalasin B. Cells were then resuspended in KHM buffer at ∼ 3 × 10^7^ cells/ml, dounce-homogenized with a tight pestle and the crude extract was centrifuged at 100,000 g for 15 min at 4°C. Latrunculin B (5 µg/ml) was added to the supernatant to decrease actin polymerization. *In vitro* microtubule aster assembly was initiated by the addition of 2.5 mM ATP and 10 µM Taxol and incubating the lysates at 30°C for 60 min. The reaction mixture was centrifuged through a 60% sucrose cushion (prepared in KHM buffer) at 100,000 g for 40 min at 4°C. Supernatant and microtubule enriched pellet fractions were analyzed by immunoblot analysis. CFDP1 immunodepleted lysates were prepared by incubating Latrunculin B supplemented mitotic extracts with Dynabeads Protein G magnetic beads pre-bound to mouse anti-CFDP1 antibody for 30 min at 4°C. After two successive rounds of immunodepletion, extracts were used for aster assembly as above in the presence or absence of full length recombinant CFDP1 protein.

### Immunofluorescence Microscopy

For 6XHIS, CENPA and β−tubulin immunofluorescence studies, cells grown on glass bottom chamber slides (Millicell EZ slide, Millipore) were fixed with cold (−20°C) ethanol containing 5% (v/v) acetic acid for 10 min and then rehydrated in cold PBS containing 0.5% Triton X-100 for 5 min. For immunofluorescence experiments with MAD1 and nocodazole wash out assays, cells were fixed with −20°C Methanol for 10 min and rehydrated as above. Blocking was performed with 1% BSA followed by incubation with primary antibodies (Table S1) diluted in PBS-0.1%Tween20 for 1 hour at room temperature. Alexa-Fluor conjugated secondary antibodies were used for detection and slides mounted with ProLong Diamond antifade reagent containing DAPI (ThermoFisher Scientific). Cells were imaged with a 40X objective on a confocal laser scanning microscope (Zeiss LSM 780). Raw immunofluorescence signals were acquired and processed with the help of ZEN application software (Zeiss) and mounted using Photoshop (Adobe).

### K-fiber length and Microtubule regrowth assays

For K-fiber length and cold-stability assay, cells were washed with PBS and incubated on ice for 15 min in L15 medium (Sigma) supplemented with 20 mM HEPES pH 7.3 followed by cold methanol fixation and immunofluorescence (as above). A minimum of 200 cells were counted for each treatment condition to quantify spindle phenotype and K-fiber length was measured using Image J (NIH).

For K-fiber length quantification in monopolar spindles, cells were treated with 50 µM Monastrol (Sigma) for 4h followed by a 10 min cold treatment and methanol fixation. Immunofluorescence analysis was performed as above. Measurements were obtained for K-fibers from at least 100 monopolar spindles for each condition using Image J. For microtubule regrowth assays cells were incubated with 3 µM Nocodazole for 3 h and washed four times with PBS and twice with medium at 37°C. Nocodazole released cells were incubated in fresh medium and fixed at indicated times with cold methanol for immunofluorescence studies. Microtubule asters were counted for more than 80 cells for each treatment condition to obtain the average number of asters per cell.

### Aurora A activation assay

Mitotic HeLa cells were collected by shake-off after 20-22 h of nocodazole treatment, washed twice with cold PBS and incubated for 5 min on ice in 0.4X EBS buffer (20 mM EGTA, 80 mM β-glycerophosphate, 100 mM sucrose, 15 mM MgCl2, 2 mM ATP, 1 mM DTT). Cells were collected and resuspended in 1X EBS supplemented with protease inhibitors and dounce-homogenized with a tight pestle. The supernatant was collected by centrifuging the homogenate at 100,000 g for 30 min at 4°C. The supernatant was further cleared by centrifugation for two more times, and collected as HeLa mitotic cytosol. For Aurora A activation assay, mitotic cytosol was incubated with 3 µM of full length or fragments of CFDP1 for 30 min at room temperature followed by addition of 20 µl of pre-assembled microtubules (5 µM). After 30 minutes of incubation, microtubule-bound proteins were separated by centrifugation at 100,000 g for 30 min at 25°C and analyzed by immunoblot assay.

### Co-Immunoprecipitation and Western blotting assays

Nuclear and cytoplasmic fractionation was carried out in NIH3T3 cells as descried previously ^[4]^. Briefly, cells were collected in ice-cold PBS and suspended in hypotonic buffer (10 mM Hepes pH8.0, 1.5 mM MgCl_2_, 10mM KCl, 0.5 mM DTT and 1x Protease inhibitors, Roche) and homogenized with a douncer to release nuclei. Homogenates were centrifuged at 2000 rpm and cytosolic fraction separated. Nuclear pellet was homogenized in Nuclear extract buffer (20 mM Hepes, pH8.0, 25% Glycerol, 1.5 mM MgCl_2_, 420 mM NaCl, 0.2 mM EDTA, 0.5 mM DTT and 1X protease inhibitors), rotated on an orbital shaker for 1 h at 4°C and nuclear extract separated by centrifugation at 13,000 rpm for 10 min at 4°C. Mitotic cytosol and mitotic chromatin extracts were prepared as above using only mitotic cells. Immunoprecipitation reactions were carried out by incubating nuclear extracts or cytosolic extracts with primary antibodies overnight at 4°C. Immune complexes were pulled down with Dynabeads Protein G magnetic beads (Invitrogen), washed thrice with wash buffer (20 mM Tris.Cl pH 8.0, 10% Glycerol, 250 mM NaCl, 5 mM EDTA, 0.1% NP40, 5 mM DTT) and bound proteins eluted in SDS-PAGE sample buffer for immunoblot analysis.

For TPX2 immunoprecipitation assays, mitotic HeLa cells were lysed in TEGN buffer (10 mM Tris pH7.4, 1 mM EDTA, 10% glycerol, 0.5% NP40, 150 mM NaCl, 1 mM DTT, 10 mM β-glycerophosphate and 1X Protease inhibitors). Pre-cleared mitotic extracts were incubated with Dynabeads Protein G magnetic beads pre-bound to mouse anti-TPX2 antibody or mouse IgG for 3h at 4°C. Beads were then washed three times with TEGN buffer and bound proteins eluted by in SDS-PAGE sample buffer for immunoblot analysis.

Whole cell lysates were prepared by incubating cell pellets with RIPA buffer (50 mM Tris.Cl pH 8.0, 1% NP40, 0.5% Sodium deoxy cholate, 150 mM NaCl, 0.1% SDS, 1 mM EDTA and 1X protease inhibitors) for 1 h at 4°C. Proteins were quantified using the BCA protein assay kit (Thermo Scientific) and equal amounts denatured in SDS-PAGE sample buffer.

Proteins for immunoblot assays were resolved in 4-20% gradient acrylamide gels (Bio Rad), transferred to PVDF membranes (Millipore) and probed with primary antibodies. Proteins were detected by the ECL method using SuperSignal West Pico PLUS Chemiluminescent Substrate Kit (Thermo Scientific).

### GST pulldown and competition assays

GST and HIS tagged proteins for bindings assays were first exchanged into binding buffer (20 mM Hepes, pH 8.0, 70 mM KCl, 10 mM MgCl2, 10% glycerol). GST-TPX2 NLS fusion protein immobilized on magnetic glutathione particles (Promega) was incubated with CFDP1 proteins in transfer buffer (20 mM Hepes pH7.4, 110 mM KCH3COO, 2mM Mg (CH3COO)2, 20% glycerol for 1 h at 4°C. Beads were washed thrice with transfer buffer and bound proteins eluted in SDS-PAGE sample buffer for immunoblot analysis. For competition assays, GST-TPX2 NLS immobilized magnetic beads were first incubated with importin α ΔIBB protein. Bead-protein complex was washed and incubated with CFDP1 proteins in transfer buffer for 1h at 4°C and processed as above for immunoblot analysis. SDS-PAGE gels were stained with QC Colloidal Coomassie stain (Bio-Rad) to visualize proteins.

### Crude chromatin preparation

Small-scale biochemical fractionation was performed to purify cytosolic, nuclear and chromatin-enriched fractions from asynchronous or mitotic NIH3T3 cells. ∼1 x 10^7^ cells were washed with cold 1X PBS and suspended in Buffer A (10 mM Hepes, pH 8.0, 10 mM KCl, 1.5 mM MgCl2, 0.34 M Sucrose, 10% Glycerol, 1 mM DTT and 1X Protease inhibitors). Triton X-100 was added to a final concentration of 0.1% and cells incubated on ice for 8 min. The cell suspension was centrifuged at 1300 g for 5 min at 4°C and the resulting nuclear pellet incubated in Buffer B (3 mM EDTA, 0.2 mM EGTA, 1 mM DTT and 1X Protease inhibitors) for 30 min on ice. Soluble chromatin was separated from the insoluble chromatin by centrifugation at 1700 g for 5 min at 4°C. The pellet fraction consisting of insoluble crude chromatin was washed once more with Buffer B and lysed with RIPA buffer for immunoblot assays.

### 3X FLAG Affinity purification

Nuclear extracts from NIH3T3 cells stably expressing 3X FLAG-CFDP1 were used for immunoprecipitation of CFDP1 interacting proteins. 500 µl (500 µg total protein) of nuclear lysate was diluted with lysis buffer (50 mM Tris.Cl pH 8.0, 1 mM EDTA and 0.5% Triton X 100) and incubated with 50 µl of FLAG M2 magnetic beads (Sigma) on a rotator at 4°C for ∼16 h. FLAG magnetic beads were washed with 20 packed column volumes of 1X TBS (50 mM Tris.Cl pH 8.0, 150 mM NaCl, 0.05% Triton X 100). Bound proteins were eluted twice in 5 packed column volumes of 1X TBS containing 3X FLAG peptide (Sigma, final concentration of 150 ng/µl). Eluates were concentrated using Amicon Ultra Centrifugal Filters (Millipore) and run on SDS-PAGE gels for Mass Spectrometry analysis as described in Supplementary materials.

### Statistical analysis

All data are presented as mean ± SD (Standard deviation) and obtained from a minimum of three experiments. *N* values are shown in respective figure legends. Data was analyzed using Microsoft Excel, GraphPad and ImageJ (NIH). An unpaired student’s *t* test was used to determine the two-tailed *P* value and considered to be statistically significant at *P* < 0.05. When significant, *P* values are mentioned in the figures.

## Acknowledgements

Generous funding by NIDCR grant DE013905 is appreciated. Confocal microscopy, MS analysis and flow cytometry were conducted at the UTSW core facilities.

## Author Contributions

GG performed the experiments, and GG, XL, and TGH wrote the manuscript.

## Data availability

All data needed to evaluate the conclusions in the paper are present in the paper and/or the Supplementary Materials.

## Figure legends for Supplementary Figures

**Fig. S1. Targeting constructs and knockout strategy for *Cfdp1* knockout and conditional knockout mice.** (**A**) Schematic detailing targeting vector construction for the generation of *Cfdp1* knockout mice. Exon 1 is replaced by a *LacZ-Neo* cassette in targeted mice. (**B**) Southern blot analysis of targeted embryonic stem cells (upper) and genotyping PCR confirmation of *Cfdp1* targeted mice (lower). (**C**) Embryonic assessment of lethality in *Cfdp1* knockout embryos. (**D**) Targeting vector design for the generation of *Cfdp1* conditional knock out mice. Exon 1 is flanked by *flox* sites and can be excised by CRE expression. (**E**) Southern Blot verification of the targeting construct and genotyping PCR confirmation of conditional knockout mice.

**Fig. S2. CFDP1 cell fractionation, localization and chromosome segregation defects in CFDP1 depleted cells.** (**A**) Immunoblot for CFDP1 levels in cytosolic, nuclear and chromatin enriched fractions prepared from NIH3T3 cells. (**B**) Immunoblot analysis for CFDP1 in G2/M phase arrested NIH3T3 mitotic chromosome fraction, corresponding supernatant and the whole cell lysate. (**C**) Immunofluorescence analysis demonstrating CFDP1 localization within the nucleus. CFDP1 (red) is targeted to distinct foci which overlap with DAPI dense foci. (**D**) Immunoblot analysis demonstrating knockdown of CFDP1 protein levels in CFDP1 siRNA treated NIH3T3 cells. (**E,F**) Chromosome segregation defects in *Cfdp1* conditional knockout Mouse embryonic fibroblasts (MEFs). Immunofluorescence analysis for tubulin in uninduced MEFs (**E**) and 4-Hydroxy Tamoxifen induced (**F**) MEFs. DNA is visualized using DAPI. (**G**) Representative immunofluorescence staining for tubulin demonstrating multi-pole spindle defect in NIH3T3 cells treated with CFDP1 siRNA.

**Fig. S3. CFDP1 interacts with components of the chromosomal passenger complex.** (**A**) CFDP1 and AuroraB immunoprecipitates from HeLa mitotic extracts were subjected to immunoblot analysis for Aurora B, INCENP, Aurora A and CFDP1 as indicated. Mitotic cells were prepared using Nocodazole as described in the main methods.

**Fig. S4. CFDP1 localization after nocodazole washout and NLS mapping of CFDP1 protein.** (**A,B**) Representative immunofluorescence analysis demonstrating CFDP1 localization at microtubule nucleation sites in NIH3T3 cells recovering from Nocodazole washout. Cells were fixed after 1 minute and stained for CFDP1 and Tubulin. DNA was visualized using DAPI. (**C**) cNLS Mapper prediction for a monopartite nuclear localization signal (NLS) in mouse CFDP1 protein. Position of NLS amino acids is highlighted (red) within the protein sequence.

## Supplementary Text

### Generation of animal models

*Cfdp1*-KO targeting vector was generated by sub-cloning a ∼9.6 Kb genomic region from a positively identified BAC clone using a homologous recombination-based technique. The short homology arm (SA) extends 1.2 Kb from the 3’ end of *Cfdp1* exon 1 while the long homology arm (LA) begins before the start codon (ATG) at the 5’ end of *Cfdp1* exon 1 and is ∼ 8 Kb long. *Cfdp1* deletion was generated by inserting a *LacZ - Neo* cassette after the start codon replacing ∼376 bp within the *Cfdp1* exon 1 coding sequence. Gene targeting of *Cfdp1* in 129/Sv – derived D3 embryonic stem cells (ES) were carried out using homologous recombination. Targeted ES colonies were identified by Southern Blot hybridization of genomic DNA digested by *PstI*. A 329 bp probe based in intron 1 was used in the hybridization assay as an external probe to distinguish the wild type band (∼ 5.1 Kb) and the mutant band (∼3.8 Kb). Chimeric mice were generated by injecting *Cfdp1* targeted ES cell lines into C57BL/6 blastocysts. Male and female *Cfdp1* KO mice were maintained as heterozygotes.

*Cfdp1* conditional KO targeting vector was generated from a 10.57 Kb genomic region sub-cloned from a BAC clone. The long homology arm (LA) extends 5.77 Kb 5’ to the location of lox P cassette inserted 1.84 Kb 5’ to the start of *Cfdp1* exon 1, while the short homology arm (SA) is 2.47 Kb in length and extends 3’ to exon 1. A *LoxP/FRT* flanked-Neo cassette was inserted 341 bp 3’ to *Cfdp1* exon 1. The target region excised upon *Cre* recombinase expression is 2349 bp long and includes the entire exon 1 coding region of *Cfdp1*. *Not I* linearized targeting vector was electroporated into BA1 (C57BL/6 × 129 SvEv) (Hybrid) ES cells. Recombinants were identified after G418 selection using PCR analysis and Southern blot analysis. Targeted hybrid ES cells were microinjected into C57BL/6 blastocysts and resulting chimeras were mated with C57BL/6 *FLP* mice for Neo Cassette removal. Both male and female *Cfdp1* conditional KO mice were maintained as heterozygotes and bred to homozygosity for mating experiments.

*Cfdp1* KO and *Cfdp1* conditional KO alleles were identified by PCR genotyping in tail lysates generated from embryos and adult mice. Genotyping of blastocysts and preimplantation embryos were performed in lysates generated by incubating the samples at 98°C for 10 min (10µl PBS diluted 1:1 with water). *Cfdp1* KO allele was identified using wildtype (WT) and Knockout (KO) primers (Table S1), which amplify a 479 bp genomic region and a 345 bp genomic region respectively. *Cfdp1* conditional knock out mice were identified using primers NDEL2 and NDEL1, which amplify a 570 bp genomic region indicating somatic *Neo* deletion and the presence of floxed exon1. Cre mediated recombination of the floxed allele was identified by a 3-primer strategy (NDEL2; NDEL1; CFDP IND F) which amplifies a 198bp genomic region indicative of a conditional knockout event for *Cfdp1* exon1.

### Blastocyst Culture

E3.5 blastocysts were obtained from crosses between mice heterozygous for *Cfdp1* KO allele and grown on gelatinized cover slips in Embryonic Stem cell media (KnockOut DMEM) without Leukemia Inhibitory Factor (LIF) at 37°C for a total of 5 days.

Blastocyst outgrowths were imaged by phase contrast microscopy or used for histology studies.

### Mitotic chromosome isolation

Cells were harvested by mitotic shake-off after Nocodazole treatment as described in main methods. After centrifugation at 500 g for 10 min, cells were washed with cold PBS and suspended in 75 mM KCl. Cells were left on ice for 20 min and spun down at 1500 g for 5 min at 4°C and homogenized in disruption buffer (10 mM Tris-HCl, pH 7.4, 120 mM KCl, 20 mM NaCl, 0.1% Triton X-100, 2 mM CaCl_2_, 1X Protease inhibitors) by passing through a 26-gauge needle five times. Cell lysates were centrifuged at 6000 rpm for 5 min and supernatant retained. Chromosome-enriched pellets were washed once more with disruption buffer and lysed in SDS-PAGE sample buffer for immunoblot analysis.

### Flow Cytometry data analysis

Flow cytometry runs were processed by FlowJo (Flowjo LLC) using a Univariate model. Events were manually gated for DNA content. After excluding debris and doublets, single cell gated population was displayed as a histogram revealing the percentage of cells in G1, S and G2/M phases using Gaussian curves to fit each cell cycle phase. Data presented were obtained from at least three independent experiments each of which were repeated in triplicates with similar results.

### Complex mixture ID analysis by LC-MS/MS method

For Mass Spectrometry analysis, denatured samples were run on a 4-20% gradient acrylamide gel. Entire protein mixture was allowed to enter the resolving part of the acrylamide gel to a distance of ∼1 cm. Gel was stained with QC Colloidal Coomassie Stain (Bio Rad) and gel fragments containing proteins were cut and subjected to complex mixture ID analysis. Samples were run on the Orbitrap Elite Mass Spectrometer utilizing a short reverse-phase LC-MS/MS method at the University of Texas Southwestern Proteomics Core. Protein identification was carried out using Proteome Discoverer 2.2.

